# Neurite Complexity and Plasticity Markers Track Synaptic Vesicle Density in the Living Human Brain: A Multimodal PET-MRI-EEG Study

**DOI:** 10.64898/2026.10.12.758444

**Authors:** Claudio Agnorelli, Joseph Peill, Danielle Kurtin, Orla Mallon, Gabriela Sawicka, Ekaterina Shatalina, Kate Godfrey, Robin Murphy, Kirran Ahmad, Lena Oestreich, Rachael Sumner, Manfredi Alberti, Matthew B Wall, Natalie Ertl, Graham Searle, Mica Clarke, Katie Zhou, Martin Osugo, Usman Sarem, Brandon Weiss, Kyle T. Greenway, Sameer Jauhar, Andrea Fagiolini, Robin Carhart-Harris, Paul M. Matthews, Eugenii A. Rabiner, David Nutt, David Erritzoe

## Abstract

Neuroplasticity, the brain’s capacity to reorganise its structure and function, is a transdiagnostic feature across psychiatric and neurodegenerative diseases and a putative mechanism of action for pharmacological interventions. [¹¹C]-UCBJ positron emission tomography (PET), which quantifies synaptic vesicle glycoprotein 2A (SV2A) density, offers the most direct *in vivo* measure of human synaptic density, but its relationship to complementary, more scalable neuroimaging measures of structural and functional plasticity remains uncharacterised. We combined [¹¹C]-UCBJ PET, diffusion weighted imaging (DWI/NODDI), resting-state functional magnetic resonance imaging (rs-fMRI), and electroencephalography (EEG)-based sensory tasks (roving mismatch negativity, MMN; visual long-term potentiation, vLTP) in 18 healthy volunteers to test the relationship between synaptic terminal SV2A density and complementary structural and functional neuroplasticity measures, and whether these relationships were reproducible following pharmacological challenge with ketamine (n = 11) or N,N-dimethyltryptamine (DMT; n = 7). There were no statistically significant differences in PET-derived SV2A binding potential (BP_ND_), DWI-derived neurite orientation dispersion (ODI) and structural weighted degree (SWD), EEG-derived MMN repetition suppression, and late-phase vLTP P2 potentiation, 1 to 8 days following DMT and ketamine exposure compared to baseline. BP_ND_ was coupled with ODI across cortical regions before and after pharmacological challenge, and this relationship was tighter within visual, attentional, and frontoparietal control brain networks. SWD showed a weaker, more spatially restricted association with BP_ND_, primarily in regions belonging to the limbic and visual brain networks, but was not replicated post-drug. No consistent association was found between BP_ND_ and functional weighted degree (FWD), derived from rs-fMRI, at baseline or after pharmacological challenge. Frontotemporal BP_ND_ was positively associated with MMN repetition suppression (r = 0.61), and occipital BP_ND_ with late-phase vLTP P2 potentiation (r = 0.54), with the MMN-BP_ND_ association replicating after pharmacological challenge (r = 0.49). These findings link synaptic terminal SV2A density in the living human brain to neurite architecture and the capacity for short-term and Hebbian-like long-term synaptic plasticity, while dissociating these relationships from generic functional connectivity and magnitude of sensory-evoked responses. These data provide cross-modal validation for DWI- and EEG-derived measures as biological indices and surrogate measures of human synaptic plasticity.

## 1. Introduction

The brain’s capacity to modify its structure and function in response to experience, defined as neuroplasticity, is central to learning, memory, and adaptive behaviour across the lifespan (*1*). At the neurobiological level, neuroplasticity manifests across two interrelated but distinct processes. Functional plasticity refers to activity-dependent changes in synaptic strength, as captured by short-term facilitation/depression and Hebbian long-term potentiation (LTP)/depression (LTD) (*2*). These are spatiotemporally situated mechanisms whereby co-active synapses are selectively strengthened or weakened through changes in receptor functioning, intracellular signalling cascades, and synaptic vesicle dynamics. On the other hand, structural plasticity encompasses the morphological correlates of these functional changes, including dendritic spine remodelling, axonal sprouting, alterations in dendritic arborisation complexity, and ultimately the formation, stabilisation, or elimination of synapses, which collectively determine the anatomical substrate on which functional plasticity operates (*3*, *4*).

Impairments across both dimensions of neuroplasticity are increasingly recognised as transdiagnostic features of neurological and psychiatric disease. In Alzheimer’s disease (AD), converging structural and functional evidence establishes synaptic pathology as a central and early mechanism of disease, preceding neuronal death and tracking cognitive decline more closely than hallmark pathologies such as amyloid-β plaques or neurofibrillary tangles (*5*). In major psychiatric disorders, converging evidence implicates altered plasticity mechanisms as a putative biomarker of psychopathology (*6*, *7*). Evidence of cortical thinning, reduced synaptic density, and impaired LTP-like mechanisms has been found in depressive disorders (*8*) and schizophrenia (*9*). This transdiagnostic profile has prompted growing interest in interventions that target neuroplasticity directly, with the common aim of restoring or enhancing plasticity capacity (*10*). Such approaches include non-invasive brain stimulation techniques and behavioural interventions, such as cognitive training. Additionally, psychedelic drugs are increasingly also described and studied as putative neuroplastogens (*3*): potential pharmacological inducers of neuroplasticity which may be related to their growing evidence as novel treatments of psychiatric illnesses. Although definitions vary, this article utilises a broad definition of psychedelics that includes both classic (serotonergic) psychedelic agents, such as psilocybin and N,N-dimethyltryptamine (DMT), as well as non-classic psychedelic agents, such as the glutamatergic drug ketamine. The rationale behind including these distinct drugs together, despite their major pharmacological differences, is that their therapeutic applications and broad neural effects – including preliminary evidence for neuroplastic effects in preclinical models – overlap considerably (*11*).

Despite this therapeutic momentum, measuring neuroplasticity reliably and non-invasively in the living human brain remains a methodological challenge that creates a substantial translational gap with the preclinical literature. In animal models, structural and functional plasticity can be characterised at the cellular and molecular level with high resolution, enabling direct quantification of synaptic number, morphology, and efficiency (*4*). In humans, equivalent measures are largely inaccessible, and the field has relied on a set of indirect proxies that each capture a partial aspect of neuroplasticity (*7*).

The introduction of [¹¹C]-UCBJ positron emission tomography (PET), which binds to synaptic vesicle glycoprotein 2A (SV2A), a transmembrane protein expressed ubiquitously in presynaptic terminals, provided the first method for the quantification of synaptic terminal density in living humans. [¹¹C]-UCBJ has been validated against post-mortem synaptophysin density, a molecular marker of synaptic density, showing a high degree of co-localisation (*12*). [¹¹C]-UCBJ has since been applied as a biomarker of synaptic dysfunction across psychiatric disorders (e.g., addiction, schizophrenia, depression, post-traumatic stress disorder), and neurodegenerative disorders (e.g., tauopathies, Huntington’s disease, α-synucleinopathies, Parkinson’s disease and frontotemporal dementia (*13*)). Furthermore, recent investigations employed longitudinal [¹¹C]-UCBJ measurements to characterise the neuroplastic effects of pharmacological interventions, such as selective serotonin reuptake inhibitors (SSRIs), ketamine, and psilocybin (*14–18*).

However, [¹¹C]-UCBJ is not a specific marker of structural or functional plasticity *per se*, since SV2A dynamics could be influenced by both increases in synaptic density and synaptic activity (*19–21*). Furthermore, as a PET method requiring arterial cannulation, cyclotron-produced radiotracers, and specialised scanner infrastructure, PET is among the most resource-intensive neuroimaging modalities available, limited to specialist research centres and small-sample experimental studies. This makes PET largely inapplicable as a diagnostic tool for real-world mental health clinical practice.

Conversely, electroencephalography (EEG) offers a scalable, low-cost, and minimally invasive readout of brain activity. EEG-based paradigms targeting cortical plasticity have been developed and deployed across several neuropsychiatric populations and pharmacological studies (*22*). Two paradigms have attracted particular attention. The auditory roving oddball paradigm utilises tones of varying frequencies to generate a mismatch negativity (MMN) event-related potential in the frontal and temporal cortex. The task captures the brain’s predictive processing by exposing the subject to an unpredictable auditory stimulus (by changing the frequency), which is then repeated to measure the brain’s adaptation to the stimulus. The EEG response evoked by the stimulus provides an index of rapid repetition suppression, reflecting synaptic adaptation, which can be interpreted as a form of short-term synaptic plasticity and/or as prediction error signalling updating within the predictive processing framework (*23*, *24*). Visual long-term potentiation (vLTP), adapted from established rodent tetanic stimulation protocols, uses high-frequency visual stimuli and captures late-phase Hebbian-like synaptic potentiation in the occipital cortex. Modulation of the visually evoked potential is therefore a product of LTP induction measurable in a non-invasive human setting (*25*). While both paradigms have demonstrated sensitivity to drug-induced modulation of plasticity and to clinical differences across diagnostic groups, they suffer from limited mechanistic validation in humans (*22*, *26*). In fact, the cellular and molecular substrates of these EEG signals remain poorly understood, and their relationship to the synaptic machinery indexed by molecular markers has not been directly established in humans.

More indirectly, non-invasive Magnetic Resonance Imaging (MRI) techniques allow the quantification of structural and functional processes in the brain through Diffusion Weighted Imaging (DWI) and resting-state functional MRI (rs-fMRI), respectively. DWI measures the degree of structural organisation via tractography and diffusion tensor imaging (DTI), and rs-fMRI measures functional brain communication via functional connectivity. Recently, Neurite Orientation Dispersion and Density Imaging (NODDI) has extended this framework by decomposing the diffusion signal into three biophysically motivated compartments: intracellular (neurite), extracellular, and free water (*27*). The orientation dispersion index (ODI) derived from this model quantifies the angular variability of dendritic and axonal processes within a voxel, providing a non-invasive surrogate of neurite branching complexity and dendritic arborisation. The biological validity of ODI has been confirmed through histological validation studies, which demonstrated that NODDI parameters correspond closely to their tissue-level counterparts (*28*). A small number of studies have examined correlations between [¹¹C]-UCBJ PET and ODI, finding moderate associations consistent with the hypothesis that higher synaptic density is associated with a more dispersed neurite configuration (*29*, *30*). Nonetheless, the evidence base for the relationship between functional connectivity, structural connectivity, ODI, and [¹¹C]-UCBJ remains limited, especially within cortical grey matter.

The present study addresses these gaps by exploiting the complementary strengths of the three neuroimaging modalities (PET/MRI/EEG) acquired within the same individuals across two multimodal imaging sessions, approximately 2-3 weeks apart, measured before and 1-8 days after a pharmacological challenge with either ketamine or DMT (*31*). We use PET-derived SV2A as a molecular reference for synaptic density against which to validate and contextualise the biological content of DWI microstructural parameters, rs-fMRI-derived connectivity, and EEG-based indices of synaptic adaptability and plasticity, with the overarching goal of establishing a cross-validated multimodal framework for the in vivo assessment of human cortical neuroplasticity. The study pursued three specific aims:

Aim 1 was to characterise the spatial relationship between PET-derived synaptic terminal SV2A density, measured via [¹¹C]-UCBJ binding potential (BP_ND_), and three MRI-derived measures: the neurite complexity index ODI derived from the DWI NODDI sequence, the structural weighted degree (SWD) derived from DWI as the sum of fibre connections between each region and the rest of the cortex, and the functional weighted degree (FWD) derived from rs-fMRI as the degree of co-activation between each region and the rest of the cortex. We hypothesised that, consistent with prior multimodal imaging evidence (*29*), higher regional synaptic terminal SV2A density ([¹¹C]-UCBJ BP_ND_) would be positively associated with greater neurite orientation dispersion (ODI), reflecting the structural complexity that co-localises with dense synaptic organisation in grey matter. Similarly, we hypothesised that synaptic density positively correlates with SWD and FWD across the cortex.

Aim 2 was to test whether PET-derived synaptic terminal SV2A density is associated with the capacity for synaptic plasticity as measured via scalp EEG, operationalised as short-term synaptic plasticity using the repetition suppression of the roving auditory mismatch negativity (MMN) and as the magnitude of Hebbian-like visual long-term potentiation (vLTP). We hypothesised that synaptic terminal SV2A density ([¹¹C]-UCBJ BP_ND_) would positively correlate with the magnitude of both MMN repetition suppression and vLTP potentiation.

Aim 3 was to determine whether the multimodal associations identified at baseline are robust to re-test and pharmacological challenge with compounds known to modulate neuroplasticity in pre-clinical models (*32*).

All analyses are replicated across two independent atlas parcellations to assess the robustness and generalisability of findings (i.e., the functional Schaefer cortical atlas (*33*) and the CIC anatomical atlas (*34*)).

The main effect of ketamine and DMT on each separate neuroimaging metric is reported in this paper but was not the primary aim of the analysis, and the results of the ketamine intervention on PET-derived SV2A have been reported elsewhere (*14*).

## 2. Methods

### 2.1. Study participants

Eighteen healthy participants (15 males and 3 females; Age 34 ± 10; Table 1) were recruited through word of mouth and public advertisements. Initial telephone screening assessed eligibility based on predetermined criteria: absence of physical or psychiatric illness, no history or current incidence of substance abuse, and no use of psychedelics for the six months before study enrolment. Participants were screened for contraindications to MRI and research PET scanning before inclusion. Informed consent was obtained from all participants before enrolment. Ethical approval was granted by the Brent Research Ethics Committee, London, UK, following the Declaration of Helsinki guidelines.

**Table 1:** Demographic, drug, and radiotracer characteristics for all participants. SD = standard deviation.

|  | All participants (N = 18) |  | Ketamine (N = 11) |  | DMT (N = 7) |  |
| --- | --- | --- | --- | --- | --- | --- |
| Age ± SD | 34 ± 10 | | 32 ± 10 | | 37 ± 11 | |
| Sex | Male = 15; Female = 3 |  | Male = 11 |  | Male = 4; Female = 3 |  |
| Psychedelic naive | N = 3 |  | N = 0 |  | N = 3 |  |
| Last psychedelic use (months ago) ± SD | 73 ± 55.6 | | 73 ± 118 | | 8 ± 4 | |
| Time of PET/MRI scan after drug (days) | 1 (N = 11), 2 (N = 1), 7 (N = 4), 8 (N = 2) |  | 1 (N = 4), 2 (N = 1), 7 (N = 4), 8 (N = 2) |  | 1 (N = 7) |  |
|  | Scan 1 | Scan 2 | Scan 1 | Scan 2 | Scan 1 | Scan 2 |
| $[^{11}\text{C}]\text{-UCBJ}$ Injected dose (MBq) ± SD | 222.51 ± 44.19 | 200.10 ± 63.50 | 231.47 ± 45.32 | 190.57 ± 70.14 | 215.88 ± 40.15 | 209.46 ± 55.63 |
| $[^{11}\text{C}]\text{-UCBJ}$ Injected mass (ng/kg) ± SD | 3.87 ± 1.36 | 3.40 ± 1.75 | 3.95 ± 1.55 | 3.57 ± 2.08 | 3.49 ± 0.94 | 2.97 ± 1.20 |

### 2.2. Study design

The data used in this analysis come from two studies investigating the effects of a single dose of ketamine (n = 11; 1 mg/kg) or DMT (n = 7; 27.5 mg) on SV2A, measured via PET [^11^C]-UCBJ in healthy volunteers. Each participant underwent two multimodal imaging sessions comprising PET, DWI/NODDI, rs-fMRI, and EEG. On each scanning day, PET scanning was performed first, followed by DWI, structural MRI, rs-fMRI, and then the EEG tasks (vLTP and MMN) (Figure 1B). Each scanning session was performed at baseline and again 2-3 weeks later (Figure 1A). One to eight days before the second scanning session, participants received a single administration of ketamine or DMT. This time window was chosen based on preclinical evidence showing maximal increases in SV2A occurring a week after psilocybin administration in pigs (*18*). No differences in [^11^C]-UCBJ binding were detected between participants scanned at day 1-2 and day 7-8 after ketamine in our previous report (*14*).

**Figure 1.**
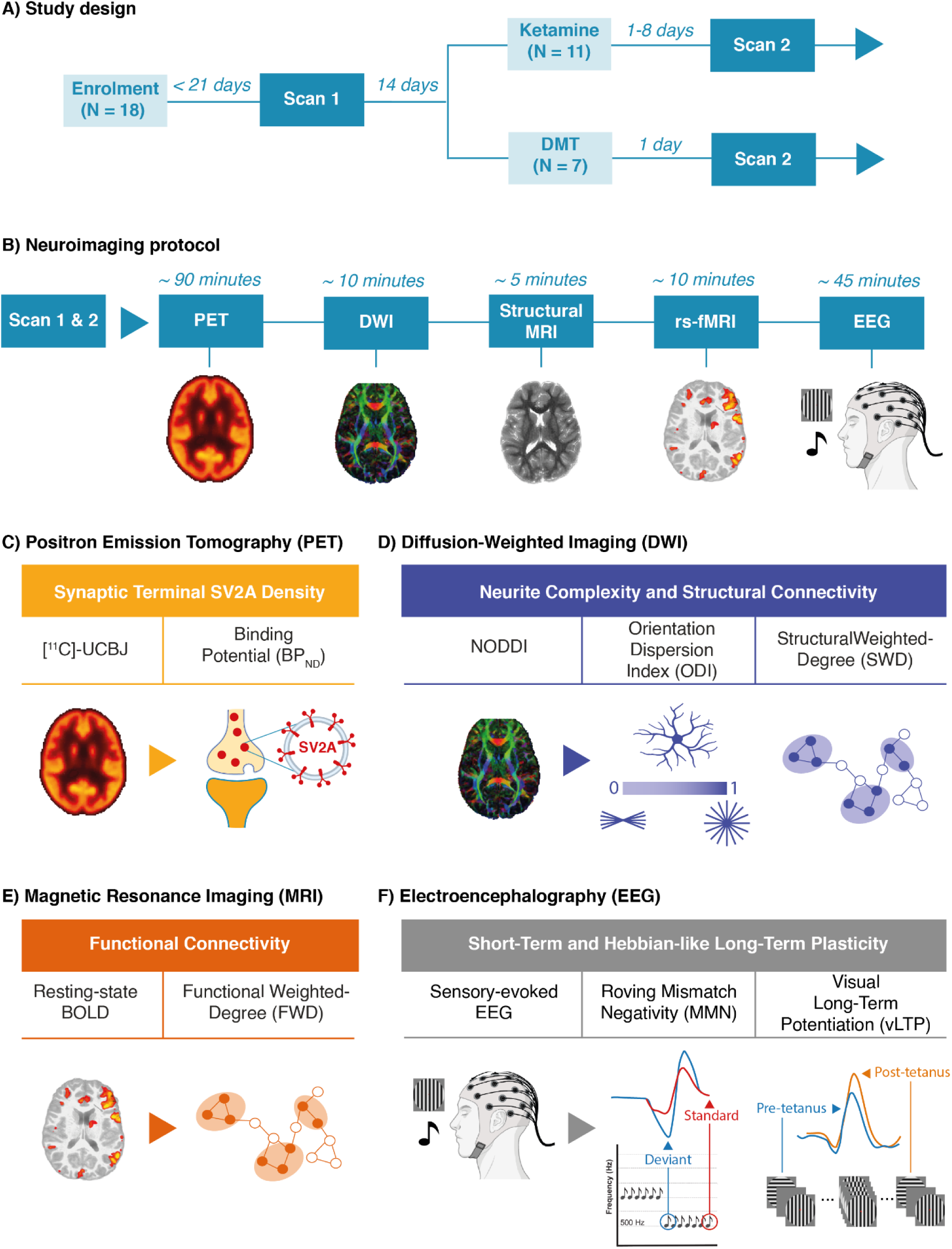
Study design and multimodal neuroimaging protocol. A) Participants (N = 18) underwent a baseline scan (Scan 1) within 21 days of enrolment and received a single dose of ketamine (n = 11) or DMT (n = 7) approximately 14 days later; the post-drug scan (Scan 2) was acquired 1-8 days after ketamine and 1 day after DMT. B) Each scanning session comprised a fixed sequence of five modalities: [¹¹C]-UCBJ PET (∼90 min), DWI (∼10 min), structural MRI (∼5 min), rs-fMRI (∼10 min) and EEG (∼45 min). C) PET was used to quantify synaptic terminal SV2A density as the non-displaceable binding potential (BP_ND_) of [¹¹C]-UCBJ. D) DWI was used to derive the NODDI orientation dispersion index (ODI), a measure of neurite complexity, and the tractography-based structural weighted degree (SWD), an index of structural connectivity strength. E) Resting-state BOLD fMRI was used to compute the functional weighted degree (FWD), an index of functional connectivity strength. F) Sensory-evoked EEG paradigms indexed short-term plasticity, as the repetition suppression of the roving mismatch negativity (MMN) from deviant to standard tones, and Hebbian-like long-term plasticity, as the potentiation of the visual evoked potential after a tetanising visual stimulus (visual long-term potentiation, vLTP).

PET, MRI, and DWI data from both studies were acquired using the same integrated General Electric Signa 3 Tesla combined PET-MRI scanner with a 32-channel head coil at the Invicro Imaging Centre (now Perceptive Discovery), London, UK. The radioligand [¹¹C]-UCBJ was synthesised onsite and administered intravenously as a bolus over 20 seconds. EEG data were acquired using the same gel-based 64-channel system (Brainvision LiveAmp 64, Brain Products GmbH). A summary of the study design and neuroimaging acquisition protocol is shown in Figure 1.

The main PET results of the ketamine study have already been published, showing no significant changes in [^11^C]-UCBJ binding 1-8 days after ketamine in 11 healthy volunteers (*14*).

### 2.3. PET acquisition and analysis

Dynamic PET data were acquired over 90 minutes following the bolus injection of [^11^C]-UCBJ. Arterial blood was sampled continuously via arterial cannulation to obtain the whole-blood activity curve, and discrete plasma samples were used to measure the plasma-over-blood ratio and the parent fraction (high-performance liquid chromatography), from which the metabolite-corrected arterial input function was derived. Images were analysed with the MIAKAT quantification tool (MATLAB, SPM12): dynamic images were corrected for motion and registered to the participant’s T1-weighted MRI, a one-tissue compartment model was fitted to estimate the total volume of distribution (V_T_), and the resulting images were registered to MNI152 space (Supplementary Methods). Regional V_T_ values were extracted for the Schaefer atlas (100 cortical parcels) (*33*) and the CIC atlas (121 cortical and subcortical grey-matter regions) (*34*). To limit partial-volume effects, regional V_T_ was averaged over grey-matter voxels only, defined as voxels with a grey-matter probability > 0.2 in each participant’s FSL FAST segmentation of the T1-weighted image, registered to MNI152 space. The non-displaceable binding potential (BP_ND_) was computed as the distribution volume ratio minus one, using the centrum semiovale (CS) as the reference region (BP_ND_ = V_T_/V_T,CS_ − 1).

### 2.4. rs-fMRI acquisition and analysis

Resting-state BOLD fMRI was acquired with an echo-planar imaging sequence (TR = 2,000 ms; TE = 30 ms; flip angle = 80°; 64 × 64 matrix; 3 × 3 mm in-plane resolution; 36 axial 3.6 mm slices; 240 volumes, 10 minutes), together with a T1-weighted IR-SPGR image (TI = 400 ms; flip angle = 11°; 1 mm isotropic voxels) used as anatomical reference. Images were converted from DICOM to NIfTI and organised in BIDS format (*35*), and preprocessed with fMRIPrep (*36*), including boundary-based registration of the T1-weighted image to the BOLD image (*37*) and the estimation of 24 motion-related confounds (translations and rotations, their temporal and spatial derivatives, and framewise displacement). The confounds were regressed out with AFNI’s 3dTproject (*38*), which, within the same linear model, applied band-pass filtering (0.01-0.20 Hz), and the data were spatially smoothed with an 8 mm FWHM Gaussian kernel.

For feature extraction, the Schaefer atlas was mapped from fsaverage to each participant’s native surface and volume with FreeSurfer (mri_surf2surf and mri_aparc2aseg). Each parcel was assigned to one of seven resting-state networks. Because no surface-based version of the CIC atlas exists, it was transformed from MNI152 space to each participant’s T1-weighted image by nonlinear registration with ANTs (affine followed by symmetric diffeomorphic normalisation) (*39*) and then resampled to the BOLD reference image; both atlases were transformed with nearest-neighbour interpolation (Supplementary Methods). The mean time series of each region was extracted with fslmeants, functional connectivity was computed as Spearman’s correlation between all pairs of regional time series, and the functional weighted degree (FWD) of each region was defined as the row-wise mean of its connectivity matrix.

### 2.5. DWI acquisition and analysis

Multi-shell DWI was acquired with an axial spin-echo EPI NODDI sequence comprising two shells (b = 1,000 and 2,000 s/mm², 60 diffusion directions each) and six b = 0 images, acquired with posterior-to-anterior phase encoding (TR/TE = 4,900/92 ms; FOV = 21 cm; matrix = 108 × 108; slice thickness = 2.5 mm), plus three b = 0 images with reversed phase encoding for distortion correction. Preprocessing combined MRtrix3 (*40*), FSL (*41*) and ANTs (*42*) and comprised denoising of each shell, Gibbs-ringing removal, susceptibility distortion correction (FSL TOPUP), eddy-current and head-motion correction with gradient-vector rotation (FSL EDDY) and N4 bias-field correction; data quality was checked with the EDDY outlier metrics. For feature extraction, fractional anisotropy (FA) and mean diffusivity (MD) were obtained by fitting the diffusion tensor to the b = 1,000 s/mm² shell (FSL DTIFIT), and ODI by fitting the NODDI model to both shells. To reduce partial-volume contamination by cerebrospinal fluid, regional NODDI metrics were computed as tissue-weighted means (*43*). The maps were registered to 2 mm MNI152 space with ANTs (rigid diffusion-to-T1 followed by nonlinear T1-to-MNI152 registration), and regional means were extracted for both atlases.

Whole-brain structural connectomes were reconstructed for each participant and session in native diffusion space following Koller et al. (*44*). Fibre orientation distributions were estimated with multi-shell multi-tissue constrained spherical deconvolution (Dhollander response functions) and intensity-normalised across tissue compartments. Anatomically constrained probabilistic tractography (iFOD2) was performed with a FreeSurfer-derived five-tissue-type segmentation, seeding from the grey-matter/white-matter interface (10 million streamlines per session; FOD amplitude cutoff = 0.06; maximum length = 250 mm), and streamlines were weighted with SIFT2 (*45*) to reduce tractography biases. Both atlases were inverse-warped into native diffusion space with nearest-neighbour interpolation, and a symmetric connectivity matrix was computed with tck2connectome, each entry being the sum of the SIFT2 weights of the streamlines connecting two regions. The structural weighted degree (SWD) of each region was defined as the row-wise sum of this matrix (for the CIC atlas, restricted to connections with the 121 grey-matter regions).

### 2.6. EEG acquisition and analysis

#### 2.6.1. Roving Mismatch Negativity (MMN) Task

The roving MMN paradigm followed the protocol described by Sumner et al. (*46*), and was presented with Psychtoolbox (*47*) in MATLAB (2023a). EEG was recorded with the 64-channel system described above. Participants passively listened via headphones, for 15 minutes, to 500 trains of 6-10 identical pure tones (70 ms, with 10 ms rise/fall ramps; stimulus onset asynchrony ∼500 ms). The tone frequency (seven frequencies between 500 and 800 Hz in 50 Hz steps) changed randomly between trains, so that the first tone of each train (S1) was a deviant that became an increasingly repeated standard over the following tones (S2-S6). To maintain alertness without directing attention to the tones, participants pressed a button when a central fixation cross briefly changed in luminance. The task was administered between the early and late post-tetanic blocks of the vLTP task (see below).

EEG data from both tasks were analysed with the FieldTrip toolbox (*48*) in MATLAB (2023a). Continuous data were band-pass filtered (0.1-30 Hz), epoched around each stimulus and baseline-corrected over the pre-stimulus interval. Noisy epochs were rejected after visual inspection, bad channels were identified visually and interpolated with spherical splines, ocular, cardiac and other non-neural components were removed with independent component analysis, and the data were re-referenced to the common average. For the MMN, epochs spanned -100 to 400 ms around each tone position (S1-S6). Based on previous literature (*49*), the MMN amplitude was quantified at Fz as the mean amplitude between 100 and 300 ms for each tone position, with the window confirmed on the grand-average waveform. MMN repetition suppression was quantified for each participant as the linear slope (first-order polynomial fit) of this amplitude across tone positions S1-S6. Because the deviant response is negative and becomes progressively less negative with repetition, a more positive slope indicates stronger repetition suppression.

#### 2.6.2. Visual Long-Term Potentiation (vLTP) Task

The vLTP task followed the protocol described by Sumner et al. (*50*) and was presented with Psychtoolbox (*47*). Participants fixated a central red dot while viewing circular sinusoidal gratings (8° of visual angle; 8 cycles per degree; ∼33 ms) on a mid-grey background, in vertical or horizontal orientation, with pseudorandomly jittered inter-stimulus intervals. The task comprised a pre-tetanus block (240 trials, both orientations at ∼1 Hz, ∼4 minutes); a tetanic block (1,000 presentations of one orientation at ∼8.4 Hz, ∼2 minutes; the tetanised orientation was counterbalanced across participants); and early (∼2 minutes after the tetanus) and late (∼30 minutes after the tetanus) post-tetanic blocks identical to the pre-tetanus block (stimulus parameters in Supplementary Methods).

vLTP data were preprocessed as described for the MMN task, with epochs from -200 to 500 ms around stimulus onset; in addition, epochs in which blinks overlapped the onset of the visual stimulus (detected semi-automatically at Fp1/Fp2 and verified visually) were rejected before independent component analysis. Event-related potentials were computed for each block by averaging the accepted epochs of the tetanised stimulus. Based on previous literature (*51*, *52*), early-phase potentiation was indexed by the N1b, measured as the mean amplitude at PO3/PO4 between 80 and 150 ms, and late-phase potentiation by the P2, measured as the peak amplitude at POz between 150 and 300 ms, with windows confirmed on the grand-average waveforms. Late-phase P2 potentiation (ΔP2-Late) was computed for each participant as the difference in P2 peak amplitude between the late post-tetanic and the pre-tetanus blocks. Because the P2 response is positive, higher ΔP2-Late reflects more potentiation.

### 2.7. Statistical analysis

Statistical analyses were performed in R (version 4.3.1) and MATLAB (2023a). Linear mixed-effects models were used for all primary inferential analyses. The primary neuroimaging outcome measures for the PET/MRI multimodal analyses were: [¹¹C]-UCBJ BP_ND_ for PET, ODI and SWD for DWI, and FWD for fMRI, extracted across cortical regions of interest. MD and FA were analysed with the same models as secondary DWI measures for completeness (Supplementary Data).

First, we fitted separate models to test for changes in each metric between the baseline and the post-drug scan. Region-level PET- and MRI-derived measures (BPND, ODI, SWD, and FWD) and the subject-level EEG-derived measures (MMN repetition suppression slope and P2 potentiation) were inserted as outcomes in the model, with Time as the fixed predictor and a random intercept for subject (e.g., BPND ∼ Time + (1 | Subject ID). The analysis of the change in each metric between the two scans was performed on the whole sample (i.e., combining the two drugs).

To test whether the different MRI-derived measures (ODI, SWD, and FWD) and EEG-derived measures are independent predictors of [¹¹C]-UCBJ BP_ND_, the degree of multicollinearity between them was assessed using the within-between mixed models described below, with either measure as predictor or outcome (e.g., SWD ∼ ODI_within_ + ODI_subject_ + ODI_region_ + (1 | Subject_ID) + (1 | ROI_Label)). The degree of multicollinearity between individually estimated MMN repetition suppression slopes and late-phase P2 potentiation (ΔP2-Late) was assessed using Pearson’s correlation. The MRI and EEG measures were largely dissociable measures of cortical structural and functional organisation and are therefore used as independent predictors throughout (Supplementary Data).

For the [¹¹C]-UCBJ BP_ND_ versus ODI/SWD/FWD relationships, each region-level predictor (ODI, SWD, or FWD) was split into a within-region component (the value minus the subject mean and the region mean, plus the grand mean) and its between-subject and between-region means, all scaled by the pooled standard deviation of the predictor. The three terms were entered as fixed effects, with ROI-level BP_ND_ as the outcome and random intercepts for subject and ROI (e.g., BP_ND_ ∼ ODI_within_ + ODI_subject_ + ODI_region_ + (1 | Subject_ID) + (1 | ROI_Label)). Each of the three terms reflects a different source of covariation between the predictor and BP_ND_. The within-region component coefficient reflects covariation across individuals within the same region, i.e., whether an individual with a higher value of the predictor than expected for that region and that individual also has higher BP_ND_; it excludes stable differences between individuals and between regions and was used for primary inference. The between-region coefficient (region-mean predictor) reflects the spatial correspondence between the group-mean maps of the two measures, i.e., whether regions with higher average values of the predictor have higher average BP_ND_ across subjects. The between-subject coefficient (subject-mean predictor) reflects whether individuals with higher overall values of the predictor across regions have higher overall BP_ND_. This within-between specification ensures that the reported within-region component slope reflects covariation across individuals within regions (*53*). Interaction terms between the within-component and atlas subdivision (cortical network or anatomical region) were included to test for regional specificity, with network-specific slopes estimated using estimated marginal trends (Kenward-Roger degrees of freedom) and corrected for multiple comparisons using the Benjamini-Hochberg false discovery rate (FDR) procedure.

For the MMN analyses, linear mixed-effect models included MMN amplitude as the outcome, tone position (S1-S6) as a fixed effect, and random intercepts for subject to characterise repetition suppression across the tone sequence (e.g., MMN ∼ Tone + (1 | Subject_ID)). For the vLTP analyses, N1b and P2 amplitude were included as the outcome, recording block (pre-tetanus, early post-tetanus, late post-tetanus) as a fixed effect, and random intercepts for subject (e.g., P2 ∼ Block + (1 | Subject_ID)). Pearson correlations were then computed between individually estimated MMN repetition suppression slopes and [¹¹C]-UCBJ BP_ND_ averaged across frontotemporal regions of interest, and between late-phase vLTP P2 potentiation (ΔP2-Late) and BP_ND_ averaged across occipital regions of interest. Regions of interest were defined a priori from the known anatomical generators of each paradigm (*49*, *51*) (frontotemporal: 22 CIC regions and 43 Schaefer parcels; occipital: 12 CIC regions and the 17 parcels of the Schaefer visual network; Supplementary Methods), and the group BP_ND_ was computed from the grey-matter volume-weighted mean V_T_ of the constituent regions.

To test whether the EEG-BP_ND_ associations were independent of neurite complexity, the EEG marker (z-scored) was added as a covariate to the ODI-BP_ND_ within-between model. For each EEG marker and atlas, the model was fitted at baseline to the ROI group used for the corresponding EEG-PET correlation (frontotemporal for MMN repetition suppression; occipital for P2 potentiation), with ODI split within the ROI group as described above: e.g., BP_ND_ ∼ ODI_within_ + ODI_subject_ + ODI_region_ + MMN + (1 | Subject_ID) + (1 | ROI_Label).

Test-retest reliability of each measure was quantified with the intraclass correlation coefficient ICC (two-way model, absolute agreement, single measure), computed on one value per subject and scan (imaging measures averaged across cortical regions of interest). The stability of the region-level associations between scans was tested with the predictor-by-scan interaction in a linear mixed model fitted to both scans (e.g., BP_ND_ ∼ (ODI_within_ + ODI_subject_ + ODI_region_) * Scan + (1 | Subject_ID/Scan) + (1 | ROI_Label)), and the stability of the subject-level EEG-PET correlations was assessed with Zou’s confidence interval for the difference between two dependent, non-overlapping correlations (*54*).

Analyses using the Schaefer atlas at baseline were considered primary; replications using the CIC atlas, V_T_, and the post-drug re-test were treated as validation and sensitivity analyses (Supplementary Data).

## 3. Results

### 3.1. Effects of ketamine and DMT on PET, MRI and EEG measures

To assess the robustness of the baseline multimodal associations, all analyses were repeated at the second timepoint in the same participants, following pharmacological challenge with drugs known to modulate neuroplasticity in pre-clinical models (11 receiving ketamine, scanned 1-8 days post-administration; 7 receiving DMT, scanned 1 day post-administration; N = 18 total). We first tested for changes in each measure in isolation after the drug challenge. We observed no statistically significant changes in BP_ND_ (consistent with the results already reported for the ketamine cohort (*14*)), ODI, SWD, FWD, MMN repetition suppression, or vLTP P2 potentiation between scan 1 (pre-drug) and scan 2 (post-drug) (see Supplementary Data for the test statistics and replication with anatomical CIC atlas and the uncorrected [^11^C]-UCBJ V_T_). These results suggest that, in our sample, DMT and ketamine did not consistently modulate synaptic terminal SV2A density, neurite complexity, structural or functional connectivity, short-term plasticity, or Hebbian-like long-term plasticity at 1-8 days after administration. No comparisons were made between the two drugs due to the small number of subjects in each group. In the following sections, each multimodal association is reported first for the baseline scan and then for the post-drug scan.

### 3.2. Relationship between synaptic terminal SV2A density and neurite complexity

At baseline, the between-region term, reflecting the spatial correspondence between the group-mean ODI and BP_ND_ brain maps across subjects (Figure 2A-B), was not significant in the Schaefer atlas, whereas it was strongly positive in the CIC atlas (β = 0.866, p < 0.001; Figure 2C). This atlas-dependent pattern reflects the wider between-region range of both measures in the volumetric CIC atlas, including a few regions with low values of both (e.g., anterior medial frontal cortex and temporal poles), and may reflect differences in tissue composition and parcellation resolution between atlases. The between-subject term, reflecting whether individuals with higher overall ODI have higher overall BP_ND_, was not significant in either atlas. Comparable between-region results were obtained at re-test and when using V_T_ (Supplementary Data).

**Figure 2.**
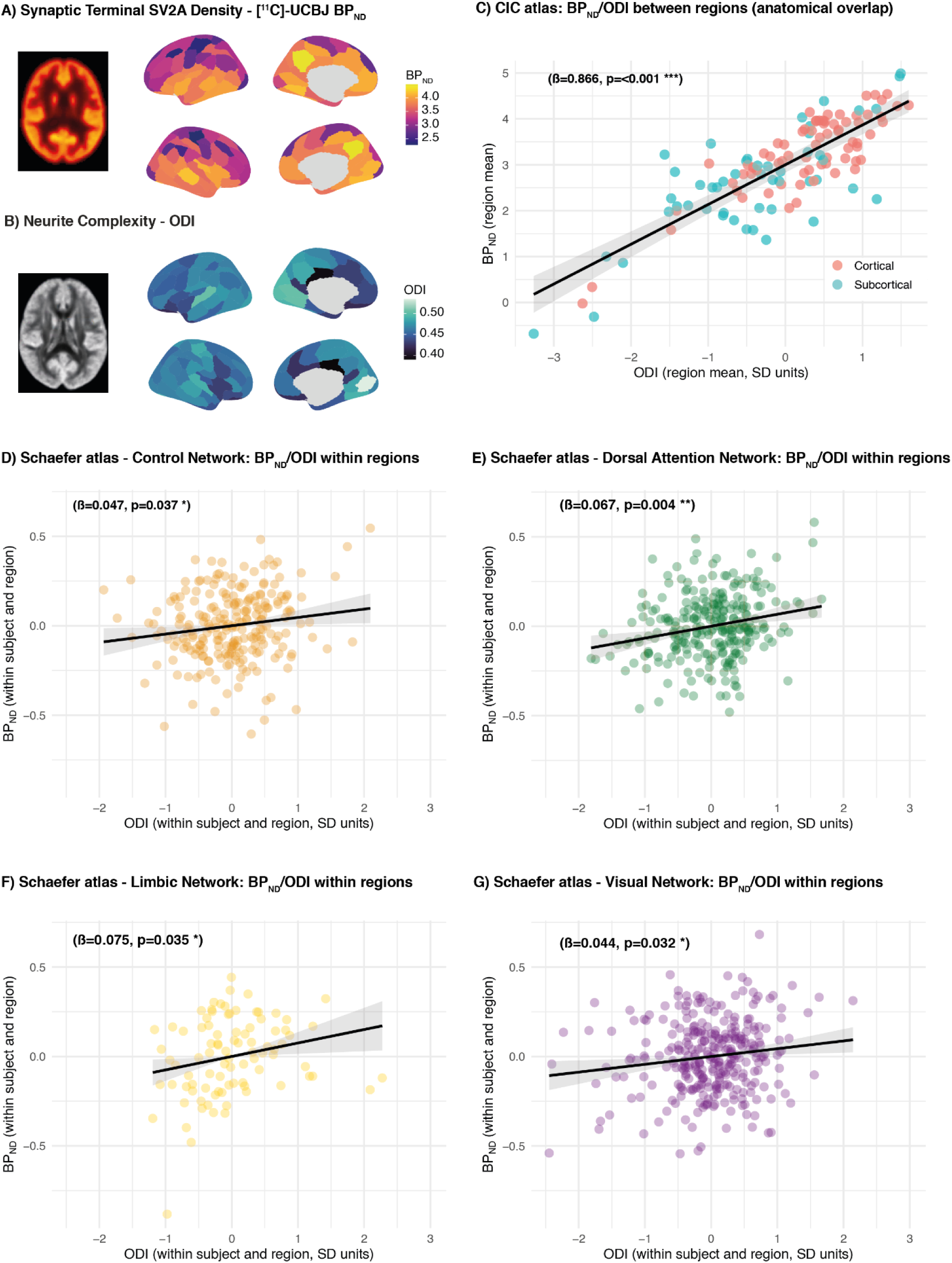
Baseline associations between synaptic terminal SV2A density and neurite complexity. A) Group-mean [¹¹C]-UCBJ BP_ND_ and B) group-mean orientation dispersion index (ODI) for the 100 parcels of the Schaefer atlas at baseline. C) Between-region association in the CIC atlas: region-mean BP_ND_ against region-mean ODI across the 121 cortical (red) and subcortical (blue) grey-matter regions. D-G) Within-region association between BP_ND_ and ODI in the Schaefer networks with a significant network-specific slope: Control (D), Dorsal Attention (E), Limbic (F) and Visual (G) networks. Each point is one subject × region observation. Predictors are expressed in units of their pooled standard deviation. Lines and shaded bands show linear fits with 95% confidence intervals. β and p-values are the corresponding coefficients of the within-between linear mixed-effects models (random intercepts for subject and region); p-values for the network-specific slopes are FDR-corrected across the seven networks. * p < 0.05, ** p < 0.01, *** p < 0.001.

The within-region term revealed a significant positive association between ODI and BP_ND_ across cortical regions (β = 0.036, p < 0.001), indicating that individuals with greater neurite complexity in a given region tend to have higher synaptic terminal SV2A density in that region, consistent with a previously reported positive association between these two neuronal markers (*29*).

Network-specific slopes revealed significant positive ODI-BP_ND_ associations in regions belonging to four brain networks: the Frontoparietal Control network (β = 0.047, p_FDR_ = 0.037; Figure 2D), the Dorsal Attention network (β = 0.067, p_FDR_ = 0.004; Figure 2E), the Limbic network (β = 0.075, p_FDR_ = 0.035; Figure 2F), and the Visual network (β = 0.044, p_FDR_ = 0.032; Figure 2G). The Somatomotor network showed a positive but non-significant slope after FDR correction. Slopes in the Default Mode and Salience/Ventral Attention networks were near zero and non-significant (Supplementary Data).

Similar results were obtained when the analysis was replicated using an anatomical atlas parcellation (CIC atlas: β = 0.139, p < 0.001), with significant positive associations across cortical (β = 0.085, p_FDR_ = 0.020) and subcortical regions (β = 0.166, p_FDR_ < 0.001). Broadly consistent results were also obtained using the uncorrected [^11^C]-UCBJ V_T_ as the PET outcome (Supplementary Data).

Following pharmacological challenge, the positive within-region association between cortical ODI and BP_ND_ was replicated (β = 0.045, p < 0.001), with significant network-specific slopes in the Visual, Dorsal Attention, Frontoparietal Control, and Somatomotor networks (Figure 5, Supplementary Data). Replication using the CIC atlas yielded a similarly significant effect (β = 0.136, p < 0.001), with significant positive slopes confirmed in both cortical (β = 0.176, p_FDR_ < 0.001) and subcortical (β = 0.118, p_FDR_ < 0.001) regions. Broadly consistent results were obtained using V_T_ as the PET outcome (Supplementary Data). The ODI-BP_ND_ slope did not differ between scans (change in β = 0.008, p = 0.459), and test-retest reliability was good for ODI (ICC = 0.86) and moderate for BP_ND_ (ICC = 0.70).

Together, these findings indicate that, especially within sensory and attentional networks, individuals with higher synaptic terminal SV2A density in a region also have higher neurite complexity in that region, and that this association is preserved after administration of ketamine and DMT.

Conventional DTI metrics were also examined for completeness. Mean diffusivity (MD) showed a consistent negative association with BP_ND_ across both atlases, both sessions, and both the within-region (Schaefer: β = -0.059, p < 0.001; CIC: β = -0.067, p < 0.001) and between-region (Schaefer: β = -0.361, p < 0.001; CIC: β = -0.551, p < 0.001) terms of the mixed models. Fractional anisotropy (FA) showed no consistent association with BP_ND_ (Supplementary Data).

### 3.3. Relationship between synaptic terminal SV2A density and structural connectivity

At baseline, the between-region term, reflecting the spatial correspondence between the group-mean SWD and BP_ND_ maps (Figure 3A-B), was negative in the Schaefer atlas (β = -0.120, p = 0.025) and not significant in the CIC atlas (β = 0.123, p = 0.188; Figure 3C), while the between-subject term was not significant in either atlas. Comparable between-region results were obtained at re-test and when using V_T_ (Supplementary Data).

**Figure 3.**
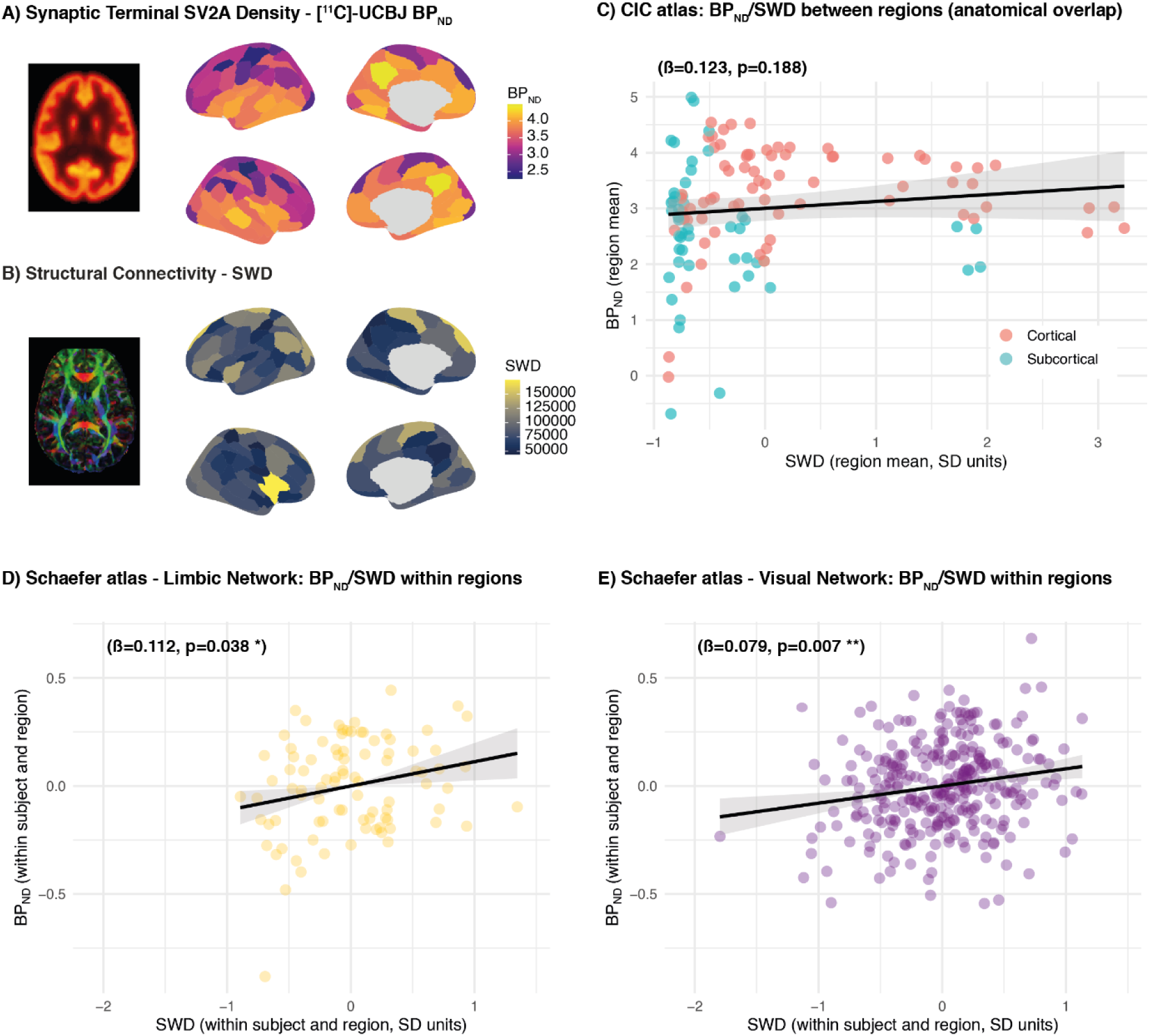
Baseline associations between synaptic terminal SV2A density and structural connectivity. A) Group-mean [¹¹C]-UCBJ BP_ND_ and B) structural weighted degree (SWD) for the 100 parcels of the Schaefer atlas at baseline. C) Between-region association in the CIC atlas: region-mean BP_ND_ against region-mean SWD across the 121 cortical (red) and subcortical (blue) grey-matter regions (not significant). D-E) Within-region association between BP_ND_ and SWD in the Schaefer networks with a significant network-specific slope: Limbic (D) and Visual (E) networks. Each point is one subject × region observation. Predictors are expressed in units of their pooled standard deviation. Lines and shaded bands show linear fits with 95% confidence intervals. β and p-values are the corresponding coefficients of the within-between linear mixed-effects models (random intercepts for subject and region); p-values for the network-specific slopes are FDR-corrected across the seven networks. * p < 0.05, ** p < 0.01, *** p < 0.001.

The within-region term showed a significant positive association between BP_ND_ and SWD across cortical regions (β = 0.026, p = 0.027). Network-specific slopes estimated via marginal trends (FDR-corrected) revealed that the SWD-BP_ND_ association was significant in regions belonging to the Limbic network (β = 0.112, p_FDR_ = 0.038; Figure 3D) and the Visual network (β = 0.079, p_FDR_ = 0.007; Figure 3E). All remaining networks showed slopes near zero and were non-significant after FDR correction (Supplementary Data). The analysis was replicated using the anatomical CIC atlas, showing a significant positive association overall (β = 0.102, *p* = 0.009). Region-specific slopes revealed a significant positive SWD-BP_ND_ association in subcortical regions (β = 0.179, p_FDR_ = 0.046), but not in cortical regions (Supplementary Data). Results using the uncorrected [^11^C]-UCBJ V_T_ as the PET outcome are reported in the Supplementary Data.

Following pharmacological challenge, the within-region SWD-BP_ND_ association across Schaefer cortical regions was not significant. Using the CIC atlas, the global association was not significant at re-test, and neither the cortical nor the subcortical slope reached significance after FDR correction (Supplementary Data). Results using V_T_ as the PET outcome are reported in the Supplementary Data.

Together, these findings suggest that the degree of structural connectivity between a given cortical region and the rest of the cortex is only weakly and inconsistently associated with its synaptic terminal SV2A density.

### 3.4. Relationship between synaptic terminal SV2A density and functional connectivity

No statistically significant within-region relationship was observed between BP_ND_ and FWD across cortical regions. Neither the between-region nor the between-subject term was significant. Using the CIC atlas, the within-region FWD-BP_ND_ association was also not significant. Results using V_T_ as the PET outcome are reported in the Supplementary Data.

Following pharmacological challenge, the within-region FWD-BP_ND_ association remained non-significant in both the Schaefer and CIC atlases (Supplementary Data). Together, these findings suggest that the degree of functional connectivity between a given cortical region and the rest of the cortex is largely unrelated to its synaptic terminal SV2A density.

### 3.5. Relationship between synaptic terminal SV2A density and short-term plasticity

Robust repetition suppression was confirmed across the tone sequence, with mean amplitude in the MMN time window increasing significantly from deviant tone to standard tone (β = 0.470, p < 0.001; Figure 4A-C). The correlation between individually estimated repetition suppression slopes and frontotemporal BP_ND_ revealed a positive association (r = 0.61, p = 0.007, N = 18; Figure 4D). Importantly, frontotemporal BP_ND_ did not correlate with the amplitude of the response to the deviant tone (r = -0.07, p = 0.779, N = 18), showing that BP_ND_ does not correlate more generally with the ERPs of prediction error sensitivity.

**Figure 4.**
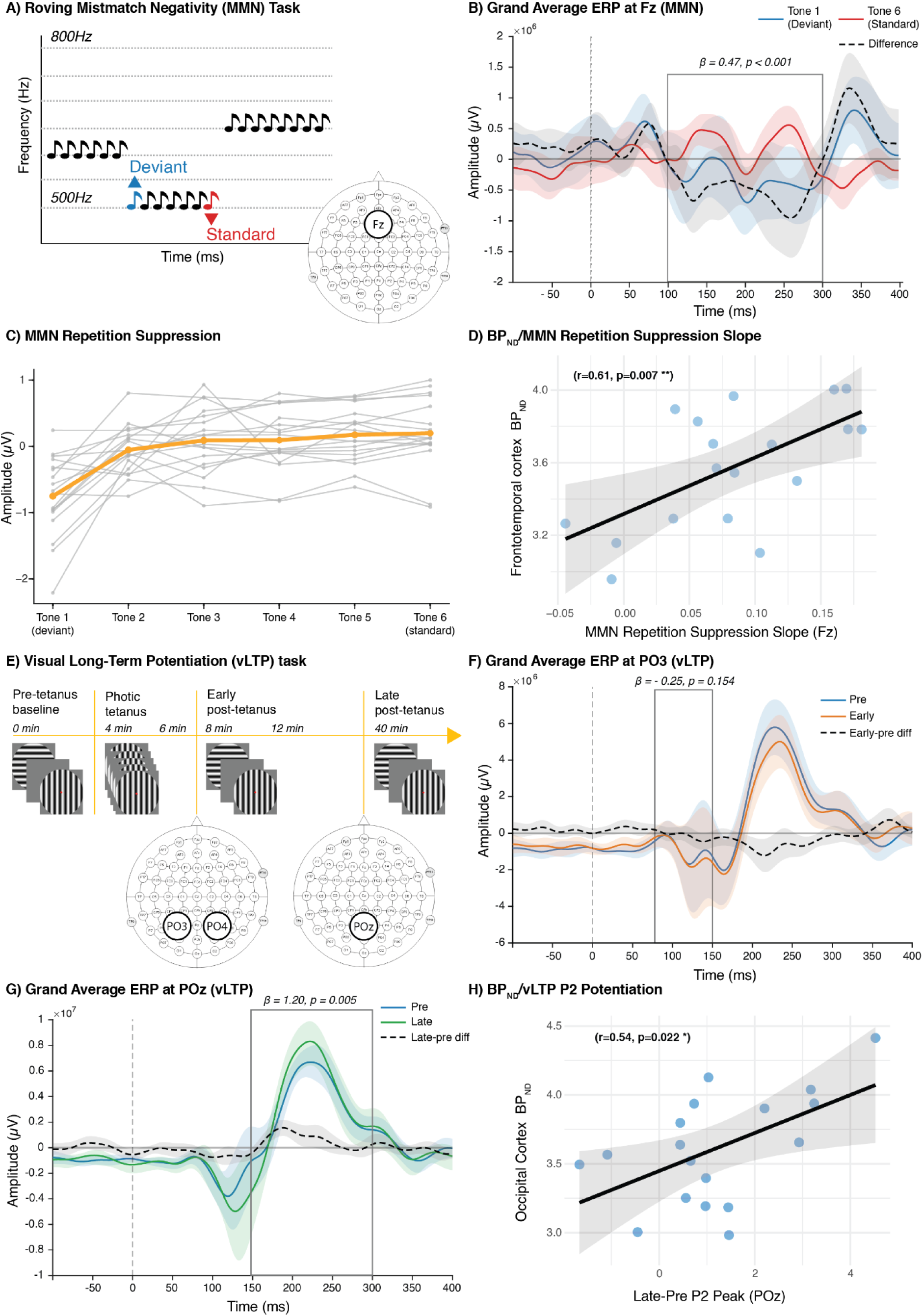
Baseline EEG markers of synaptic plasticity and their association with synaptic terminal SV2A density. A) Roving MMN paradigm: the tone frequency (500-800 Hz) changed between trains of identical tones, so that the first tone of each train (Tone 1, deviant) became a repeated standard over the following tones (Tone 6, standard). The analysis electrode (Fz) is highlighted. B) Grand-average event-related potentials (ERPs) at Fz for Tone 1 (deviant; blue) and Tone 6 (standard; red), and their difference (dashed black); the grey box marks the MMN analysis window, and β and p refer to the mixed-model comparison of Tone 6 vs Tone 1 amplitude. C) MMN amplitude at Fz across Tones 1-6 for each participant (grey) and the group mean (orange), showing the increase in amplitude from deviant to standard that reflects repetition suppression. Because the deviant response is negative and becomes progressively less negative with repetition, a more positive slope indicates stronger repetition suppression. D) Pearson’s correlation between the individual MMN repetition suppression slope and frontotemporal BP_ND_ (Schaefer atlas). E) vLTP task timeline (pre-tetanus baseline, photic tetanus, early and late post-tetanus blocks) and analysis electrodes for the N1b (PO3/PO4) and P2 (POz). F) Grand-average ERPs for the pre-tetanus (blue) and early post-tetanus (orange) blocks and their difference (dashed black); the grey box marks the N1b analysis window, and β and p refer to the mixed-model comparison of early vs pre-tetanus N1b amplitude. G) Grand-average ERPs at POz for the pre-tetanus (blue) and late post-tetanus (green) blocks and their difference (dashed black); the grey box marks the P2 analysis window, and β and p refer to the mixed-model comparison of late vs pre-tetanus P2 peak amplitude. H) Pearson’s correlation between late-phase P2 potentiation (late minus pre-tetanus P2 peak amplitude at POz) and occipital BP_ND_ (Schaefer atlas). Because the P2 response is positive, higher ΔP2-Late reflects more potentiation. Shaded areas around the ERPs indicate between-participant variability; lines and shaded bands in D and H show linear fits with 95% confidence intervals. N = 18.

Similar results were obtained using the anatomical CIC atlas (r = 0.61, p = 0.007, N = 18). However, the result was not replicated with the uncorrected [^11^C]-UCBJ V_T_ metric (Supplementary Data).

Following pharmacological challenge, the MMN effect was replicated (β = 0.529, p < 0.001). Further, frontotemporal BP_ND_ and the MMN repetition suppression response remained positively correlated in the post-drug scan, observed using both the Schaefer atlas (r = 0.49, p = 0.038; Figure 5) and the CIC atlas (r = 0.52, p = 0.026). The direction and magnitude of the correlation were consistent with the baseline estimate (r = 0.61). Notably, V_T_ showed a significant positive correlation with MMN repetition suppression at re-test in both the CIC (r = 0.61, p = 0.007) and Schaefer atlases (r = 0.60, p = 0.009) (see Supplementary Data). The MMN repetition suppression-BP_ND_ correlation did not differ significantly between scans (Δr = -0.12, 95% CI -0.57 to 0.30), and test-retest reliability was moderate for both MMN repetition suppression (ICC = 0.56) and frontotemporal BP_ND_ (ICC = 0.71).

**Figure 5.**
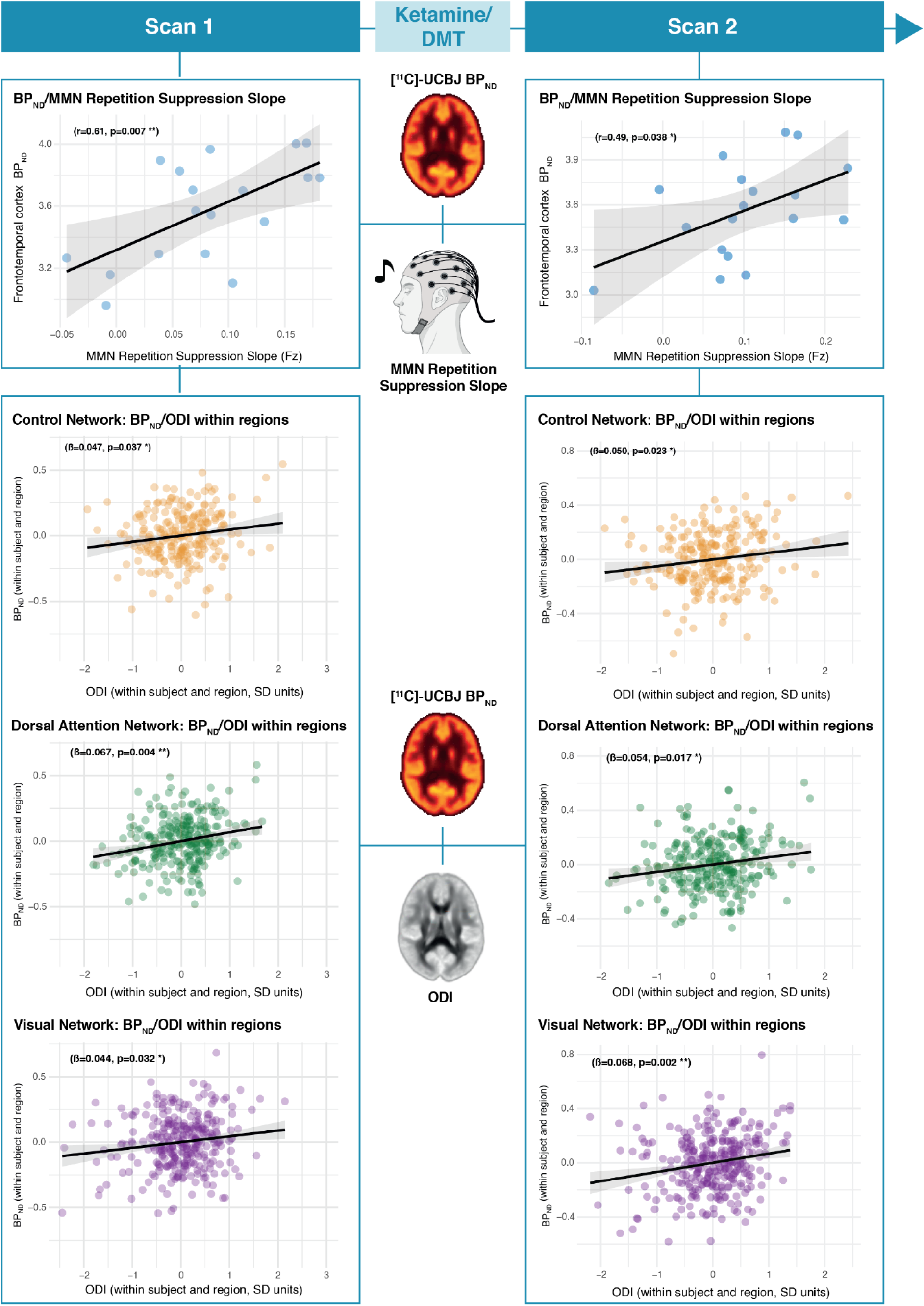
Correlations with synaptic terminal SV2A density that were significant at both baseline (Scan 1) and post-drug re-test (Scan 2) (Schaefer atlas). Left column: baseline; right column: re-test after a single dose of ketamine or DMT. Top row: Pearson’s correlation between the individual MMN repetition suppression slope at Fz and frontotemporal BP_ND_. Lower rows: within-region association between BP_ND_ and ODI in the Control, Dorsal Attention and Visual networks. * p < 0.05, ** p < 0.01.

These findings suggest that frontotemporal synaptic terminal SV2A density correlates with the capacity for short-term repetition-based synaptic modification underlying adaptation to auditory stimuli.

### 3.6. Relationship between synaptic terminal SV2A density and Hebbian-like long-term plasticity

The vLTP N1b mean amplitude change from pre to early post-tetanus was not significant (β = −0.248, p = 0.154; Figure 4E-F). This indicated that N1b was not potentiated by the tetanus stimulation. Therefore, no correlation test was performed between N1b and BP_ND_.

P2 peak amplitude was significantly larger at the late post-tetanus condition relative to the pre-tetanus block (β = 1.201, p = 0.005; Figure 4G), confirming robust late-phase visual LTP. There was a statistically significant positive correlation between ΔP2-Late (late minus pre-tetanus P2 peak amplitude) and occipital BP_ND_ (r = 0.54, p = 0.022, N = 18; Figure 4H) at baseline. There was no correlation between pre-tetanus P2 peak amplitude and occipital BP_ND_ (r = 0.11, p = 0.676, N = 18), confirming no relationship between BP_ND_ and general visual ERP amplitude. Similar results were obtained using the anatomical CIC atlas and using the uncorrected [^11^C]-UCBJ V_T_ as the PET outcome (Supplementary Data).

Following pharmacological challenge, the late-phase P2 potentiation effect observed at baseline was not reproduced (β = 0.580, p = 0.326), which may indicate either a trending effect of the drug on vLTP or a failure to replicate the vLTP effect on the visual ERP. Because this potentiation effect was absent, we could not meaningfully evaluate the correlation between P2 potentiation and occipital BP_ND_ in this dataset.

These findings suggest that occipital synaptic terminal SV2A density scales with the capacity for LTP-like plasticity in the visual cortex, consistent with a role for presynaptic vesicle machinery in sustaining the long-term strengthening of synaptic connections following high-frequency stimulation.

### 3.7. Associations between EEG plasticity markers and synaptic terminal SV2A density after adjustment for neurite complexity

To test whether the EEG-BP_ND_ associations were independent of neurite complexity, the EEG marker was added as a covariate to the ODI-BP_ND_ within-between model, fitted at baseline within the ROI group used for each EEG-BP_ND_ association. In the Schaefer atlas, MMN repetition suppression remained associated with frontotemporal BP_ND_ after adjustment for ODI (β = 0.210, p = 0.003), and late-phase P2 potentiation with occipital BP_ND_ (β = 0.199, p = 0.023). Similar results were obtained with the CIC atlas (MMN repetition suppression: β = 0.186, p < 0.001; P2 potentiation: β = 0.238, p = 0.010; Supplementary Data).

Together, these findings indicate that the associations between synaptic terminal SV2A density and the EEG plasticity markers are not explained by neurite complexity in the same regions.

## 4. Discussion

The present study employed an innovative multimodal neuroimaging approach, combining [¹¹C]-UCBJ PET, DWI, fMRI, and EEG plasticity paradigms (roving MMN and vLTP) within the same individuals, to cross-correlate and validate complementary measures of synaptic microstructure and plasticity capacity with molecular imaging of synaptic terminal density in the living human brain, and to test the impact of pharmacological challenge with ketamine and DMT on these associations.

First, we observed that neurite complexity (ODI) was robustly and positively associated with synaptic terminal SV2A density (BP_ND_) within cortical regions, with the association concentrated in the Visual, Dorsal Attention, and Control networks, and this association was consistently replicated at re-test following pharmacological challenge. Second, the degree of structural connectivity of a given region with the rest of the cortex (SWD) showed a weaker and more spatially restricted relationship with synaptic terminal SV2A density, with significant effects primarily in visual and limbic regions, but it was not replicated following pharmacological challenge. Third, EEG indices of short-term plasticity (MMN repetition suppression slope) and Hebbian-like long-term plasticity (late-phase vLTP P2 potentiation) were positively correlated with regional synaptic terminal SV2A density at the individual level, with the MMN repetition suppression-BP_ND_ association replicating at post-drug re-test. Taken together, these findings reveal that [^11^C]-UCBJ BP_ND_ may reflect a composite signal of both structural and functional synaptic determinants. Our multimodal approach also enabled cross-modal validation between DWI and EEG with PET, revealing relationships between neurite complexity, synaptic terminal SV2A density, and functional plasticity measures.

The positive association between cortical ODI and [¹¹C]-UCBJ BP_ND_ is consistent with the anatomical logic underlying both measures. ODI quantifies the angular variability of dendritic and axonal processes within a voxel (i.e., neurite complexity), while [¹¹C]-UCBJ BP_ND_ measures synaptic terminal SV2A density, with the density directly determined by the arborisation complexity of the dendritic tree that hosts them (*55*). An individual with more complex, multidirectional dendritic branching in a given region, reflected as higher ODI, should therefore harbour more synaptic contacts per unit volume in that region. This is precisely what the within-region term of our models captures. Because this association is estimated after removing each region’s and each individual’s mean, it cannot be explained by fixed regional differences in cytoarchitecture or tissue composition, nor by global individual differences, and it was present in both atlases and at both sessions. This interpretation aligns with work by Whiteside et al., which reported a similar positive association between cortical ODI and [¹¹C]-UCBJ BP_ND_ (*29*). We extend these findings by showing a cortical network specificity of the association, whereby significant ODI-BP_ND_ coupling was observed in regions belonging to the Visual, Dorsal Attention, and Frontoparietal Control networks, but was absent in the Default Mode and Salience networks. Primary sensory regions have a more differentiated, laminated architecture, in which local neurite geometry and synaptic terminal density are closely aligned. Transmodal cortices are less laminated, and their microstructural and functional organisation are increasingly dissociated (*56–59*). In sensory and limbic regions, ODI and BP_ND_ are therefore expected to co-vary because both reflect the same underlying anatomical substrate. Executive and attention networks, including the Frontoparietal Control and Dorsal Attention networks, similarly exhibit relatively well-laminated cytoarchitecture. In contrast, the Default Mode and Salience networks, despite being among the highest [¹¹C]-UCBJ BP_ND_ regions in the brain, appear to be characterised by a partial dissociation between dendritic complexity and synaptic terminal density (*60*). This may reflect the fact that in these networks, synaptic density is more strongly determined by long-range axonal inputs, whose terminal boutons contribute substantially to local SV2A binding, but whose fibres of passage do not meaningfully influence the local ODI signal (*61*). In other words, in regions where synaptic input is heavily dominated by extrinsic afferents rather than local dendritic elaboration, the geometric relationship between neurite complexity and presynaptic terminal density might break down. The stability of the coupling between neurite complexity and synaptic terminal SV2A density across the post-drug re-test further supports the interpretation that this coupling might reflect a stable organisational property of cortical tissue rather than a state-dependent signal, and that neither ketamine nor DMT at the doses administered altered the structural microarchitecture of the cortex sufficiently to disrupt this relationship across the 1-8 day post-drug window.

The between-region term, reflecting the spatial correspondence between the group-mean ODI and BP_ND_ maps, was absent across the 100 cortical Schaefer functional parcels but highly significant across the 121 subcortical and cortical CIC anatomical parcels. The strong between-region association in the CIC atlas reflects the broader range of tissue types it covers, including subcortical nuclei and a few cortical regions with low values of both measures (e.g., anterior medial frontal cortex and temporal poles). It is therefore more likely to capture coarse differences in tissue composition between region types than a graded coupling between neurite complexity and synaptic density. Similarly, the between-subject term was not significant in either atlas: individuals with higher average ODI across the brain did not show higher average BP_ND_. With 18 participants, this does not exclude a modest global association, but it indicates that the ODI-BP_ND_ coupling is expressed mainly as local, region-specific co-variation across individuals rather than as a brain-wide individual trait.

The within-region association between structural weighted degree (SWD) and synaptic terminal SV2A density was weaker and more spatially restricted than the ODI-BP_ND_ coupling, reaching significance primarily in Limbic and Visual network regions under the Schaefer parcellation and in subcortical but not cortical regions under the CIC atlas. This pattern suggests that the coupling between axonal wiring density and synaptic terminal SV2A availability is not a global property of the brain but might instead be concentrated within specific cortical regions. The SWD metric used here, the SIFT2-weighted sum of incoming connection strengths per region, was found to exhibit a systematic spatial gradient across the human cortex, increasing from temporal and parietal towards frontal and occipital areas, and to decay steeply with inter-regional distance, such that short-range connections dominate SWD in most areas (*44*). Consistent with this, our results suggest that SWD would couple with synaptic terminal SV2A density only in regions where connectivity is dominated by short-range connectivity, such as those belonging to the Visual network (*59*). However, the small global effect size, which was not replicated at re-test in either atlas, limits the generalisability of this interpretation.

No relationship was observed between functional weighted degree (FWD) and synaptic terminal SV2A density within cortical regions, either at baseline or following pharmacological challenge with ketamine and DMT. In their multimodal study, Whiteside and colleagues reported a significant positive association between resting-state FWD and [¹¹C]-UCBJ BP_ND_ across cortical regions in patients with frontotemporal lobar degeneration syndromes. However, this same study found no such relationship in their healthy control group, demonstrating that the coupling between synaptic density and functional connectivity is not a baseline property of the healthy cortex, but may emerge specifically in the context of pathological synaptic loss (*29*). Our findings, obtained in healthy volunteers, are therefore consistent with the results of Whiteside et al., and extend them by showing that this lack of effect is also robust to short-term pharmacological perturbation with psychedelic drugs. These findings suggest that in the intact human brain, a region’s degree of functional integration with the rest of the cortex is largely dissociable from its local synaptic terminal SV2A density and that the coupling between SV2A availability and non-invasive proxy measures in the healthy brain is selective to indices more directly associated with neuroplastic processes, such as microstructural properties and functional short-term and Hebbian-like plasticity.

The positive correlation between frontotemporal [¹¹C]-UCBJ BP_ND_ and MMN repetition suppression slope, and between occipital BP_ND_ and late-phase vLTP P2 potentiation, provides the first direct *in vivo* evidence linking synaptic terminal SV2A density to the functional capacity for plasticity in the human brain. The MMN repetition suppression index, defined as the parametric rate of amplitude change across tone positions S1-S6 in a roving-standard MMN paradigm, reflects the progressive attenuation of auditory prediction error responses through stimulus-specific adaptation at auditory cortical synapses (*62*, *63*). At the cellular level, this adaptation is understood to involve a combination of network-level synaptic depression and short-term plasticity at glutamatergic synapses, and has been formally modelled as requiring both adaptation and synaptic plasticity components to fully account for the roving-standard MMN effect (*49*, *64*). The finding that individuals with higher frontotemporal synaptic terminal SV2A density show greater rates of repetition suppression suggests that higher density of SV2A-expressing presynaptic terminals provides a larger substrate for short-term activity-dependent modification, thereby enabling more rapid and pronounced amplitude adaptation across tone repetitions. Critically, BP_ND_ was associated with the rate of adaptation rather than with the amplitude of the deviant ERP response. This confirms that synaptic terminal SV2A density does not simply scale the magnitude of auditory-evoked responses but specifically indexes the capacity for dynamic, plasticity-related, synaptic adaptations. The replication of the MMN repetition suppression-BP_ND_ coupling at re-test, in the context of pharmacological manipulation by ketamine and DMT, across anatomical and functional brain parcellations, supports the reliability of this association as a stable trait.

The vLTP result extends this interpretation to the domain of Hebbian-like long-term plasticity. Occipital BP_ND_ was associated with the magnitude of late-phase P2 potentiation following high-frequency tetanic stimulation, a paradigm whose LTP-like characteristics, including NMDA receptor dependence, input specificity, stimulus orientation selectivity, and persistence over at least 30 minutes, have been established in prior human EEG studies (*25*, *50*, *65*). That individuals with greater occipital synaptic terminal SV2A density show larger late-phase potentiation suggests that a denser presynaptic terminal population supports the sustained vesicle release necessary to drive and maintain postsynaptic AMPA receptor insertion during LTP consolidation. Like the MMN repetition suppression-BP_ND_ coupling, no relationship was found between the magnitude of the P2 evoked potential itself and occipital BP_ND_, showing that the coupling was specific to the P2 potentiation associated with long-term Hebbian-like plasticity. No significant N1b potentiation was observed in our sample, which precluded a meaningful test of whether the early post-tetanic component modulation also relates to occipital synaptic density. Whether the absence of N1b potentiation reflects a genuine ceiling or floor effect in our healthy volunteer sample, differences in task compliance or attention during the early post-tetanic block, or simply limited power at this effect size, remains an open question for future studies with larger samples. Of note, the potentiation of the N1b component is not always reliably found with the protocol as used (*51*). The failure to replicate late-phase P2 potentiation following the pharmacological challenge meant that the stability of the P2-BP_ND_ coupling across sessions could not be evaluated. This represents an important limitation relative to the MMN finding, where test-retest stability of the PET-EEG association was directly demonstrated. The absence of P2 potentiation post-drug in our sample may speak to complex interactions between the two different drugs (ketamine and DMT), repetition effects on the task, and/or lack of power in the ability to separate the ERPs themselves. Larger longitudinal designs, incorporating drug-naïve re-test sessions, will be necessary to establish the reliability of the vLTP-synaptic density relationship.

Considered together, a central implication of our findings is that [¹¹C]-UCBJ BP_ND_ does not reflect a single, unitary biological process but rather a composite signal with both structural and functional neuroplastic determinants. Mechanistically, SV2A availability tracks synaptophysin and synapse density in post-mortem tissue (*12*), but SV2 also primes docked vesicles and increases release probability during low-frequency activity (*20*). On one hand, the robust ODI-BP_ND_ association suggests that a substantial portion of inter-regional and inter-individual variance in synaptic terminal SV2A density is explained by the structural complexity of the dendritic arbour, a relatively stable anatomical feature. On the other hand, the positive correlations with both MMN repetition suppression and vLTP demonstrate that BP_ND_ coupling also scales with the functional capacity for activity-dependent synaptic modification, a functional, experience-dependent property. Importantly, the relationship between [¹¹C]-UCBJ BP_ND_ and the EEG functional markers survived adjustment for ODI, and these markers did not correlate with each other. This dual sensitivity is consistent with SV2A’s molecular biology, being a protein expressed in the presynaptic terminal of virtually all neurons, irrespective of their functional state. This interpretation is further supported by prior work showing that [¹¹C]-UCBJ binding correlates with [¹⁸F]-FDG-PET markers of regional glucose metabolism (*66*) and the amplitude of low-frequency fluctuations (ALFF) measured via fMRI (*14*, *21*), both proxies of ongoing neuronal and synaptic activity, indicating that SV2A availability is sensitive to local activity-dependent processes. Importantly, the present findings extend this observation by showing that the functional plasticity dimension of the SV2A signal, indexed here by EEG, is anatomically specific, replicable at the individual level, and stable after a pharmacological perturbation. Notably, our finding of null associations between resting-state functional connectivity (FWD), MMN ERP, and P2 ERP with synaptic terminal SV2A density suggests that SV2A availability appears selectively coupled to structural and functional plasticity markers, while remaining dissociable from resting-state functional integration or the magnitude of sensory-evoked responses.

A secondary but important contribution of this work is the cross-modal validation it provides for NODDI and EEG as practical neuroplasticity biomarkers. Both modalities are considerably more scalable, affordable, and clinically deployable than PET, yet their biological validity as indices of synaptic processes in humans has until now rested primarily on histological validation of model parameters or on indirect associations with cognitive measures (*28*, *67–69*). ODI and the EEG plasticity indices were each associated with BP_ND_ separately. This matters because histological validation of ODI has so far come mainly from white matter and spinal cord (*28*, *68*), leaving its interpretation in cortical grey matter largely inferential. Similarly, vLTP magnitude and MMN repetition suppression slopes have not been validated against a molecular synaptic marker. By demonstrating that ODI and EEG plasticity indices are each independently associated with grey matter [¹¹C]-UCBJ BP_ND_, the closest available molecular reference standard for *in vivo* synaptic density, the present study provides a direct empirical basis for their interpretation as biologically grounded surrogates of cortical microstructure and capacity for synaptic plasticity, respectively. The stability of the ODI-BP_ND_ and MMN repetition suppression-BP_ND_ associations at re-test, in the context of pharmacological manipulation designed to modulate neuroplasticity, further supports their reliability as indices that are sensitive to genuine inter-individual biological differences. These results suggest that ODI extracted from NODDI acquisition and MMN repetition suppression from a passive EEG paradigm may offer valid and complementary proxies of synaptic plasticity.

Several limitations should be considered when interpreting these findings. The sample size of N = 18, while adequate to detect the moderate-to-large correlations observed, limits statistical power for detecting smaller effects and precludes conclusions about sex differences or drug-specific effects. The fixed-order design might have introduced order and learning effects. The two drug arms (ketamine, n = 11; DMT, n = 7) were pooled for re-test analyses given the absence of significant group-level changes in any neuroimaging metric, but drug-specific modulation of the multimodal associations cannot be excluded. The absence of vLTP replication at re-test, potentially reflecting limited power and a complex interaction with the two drugs, represents a notable limitation for that specific analysis. Finally, the cross-sectional and correlational nature of the baseline associations precludes causal interpretation.

In conclusion, these findings carry several practical implications for the design and interpretation of neuroplasticity research. For studies using [¹¹C]-UCBJ PET to track neuroplastic effects of pharmacological interventions, such as psychedelic therapies, and non-invasive brain stimulation techniques, the present results caution against interpreting changes in SV2A binding as exclusively reflecting either structural synaptogenesis or functional synaptic upregulation in isolation. A drug that increases synaptic terminal SV2A density following a single dose may do so through rapid functional upregulation of presynaptic vesicle pools, through structural spinogenesis and synaptogenesis, or through both, and the relative contribution of each process will have different implications for the timescale, durability, and therapeutic relevance of the observed effect (*3*). At the same time, the present study validates complementary non-invasive measures of neuroplasticity, suggesting that DWI-derived indices of neurite complexity and structural connectivity, and EEG-based markers of functional plasticity, co-vary with synaptic terminal SV2A density as measured by [¹¹C]-UCBJ PET. Taken together, these findings demonstrate the value of multimodal neuroimaging for dissecting the biological substrates of synaptic plasticity in the living human brain, and establish a framework against which drug-induced neuroplastic effects can be more precisely characterised.

## 5. Data availability

Processed data and code will be made open access upon publication. Address correspondence to Claudio Agnorelli at for raw data availability.

## 6. Author contributions

C.A. contributed to study design and setup, data collection, data analysis, figures, and manuscript writing. J.P. contributed to the study setup, data collection, data analysis, and manuscript writing. D.K. contributed to the data analysis and manuscript writing. O.M. contributed to data collection and manuscript writing. G.Sa. contributed to the data collection and manuscript writing. E.S. contributed to the data analysis and manuscript writing. K.G. contributed to the study setup, data collection, data analysis, and manuscript writing. R.M. contributed to data analysis and manuscript writing. K.A. contributed to the study design and setup, and data collection. L.O. contributed to data analysis and manuscript writing. R.S. contributed to the study setup and manuscript writing. M.A. contributed to data analysis. M.B.W. contributed to the study setup and manuscript writing. N.E. contributed to data analysis. G.Se. contributed to data analysis. C.M. contributed to data analysis. K.Z. contributed to the data collection. M.O. contributed to the data collection. U.S. contributed to data collection. B.W. contributed to manuscript writing. K.T.G. contributed to manuscript writing. S.J. contributed to manuscript writing. A.F. contributed to manuscript writing. R.C-H. contributed to manuscript writing. D) N. and P.M.M. contributed to the acquisition of funding, study design and setup, and manuscript writing. E.A.R. contributed to study design and setup, data analysis, and manuscript writing. D.E. contributed to the acquisition of funding, study design and setup, data collection, study supervision, and manuscript writing.

## 7. Funding

The ketamine study (n=11) was funded by GE Healthcare Global Research Organisation’s Investigator-Initiated Study Program. The DMT study (n=7) was funded by the non-profit association Drug Science.

## 8. Conflict of Interest

The primary employer of E.A.R., M.B.W., C.R. and N.E. is Perceptive Inc., a contract research organisation, which provides services to the pharmaceutical and biotechnology industries. M.B.W. has also received travel support and honoraria from Compass Pathways. D.E. acts as a paid scientific advisor for Aya Biosciences, Lophora ApS, Clerkenwell Health, Mindstate Design Lab. P.M.M. is a consultant for Biogen, Novartis, Nodthera, Sudo Therapeutics and GSK. He is receiving research funding from Perceptive and Biogen. R.C-H. is an advisor to Osmind, Otsuka, Entropy Neurodynamics, and Atai-Beckley.

D.N., via the Imperial College Centre for Psychedelic Research, has received research support from COMPASS Pathways, Usona, Filament Health, Beckley Psytech, and MAPS for psychedelic drug studies. He is a scientific advisor to Algernon and Neural Therapeutics, companies that are researching psychedelic medicines and also scientific chair of the charity Drug Science. All the other authors have no conflict of interest to disclose.

## 9. Acknowledgements

We thank GE and Drug Science for funding and support, and Imperial College London for sponsorship and study governance. We thank Central & North West London NHS Foundation Trust and its staff for supporting and hosting the ketamine study in the CNWL-Imperial collaborative research space, CIPPRes Clinic at St Charles Hospital. We thank the NOCLOR team for CIPPRes site governance. We thank Robin Tyacke, Tf Chan and Allan Sebti, for invaluable help with study drug processes, as well as Trust Directors Drs Jo Emmanual, Jasna Munjiza, and Cornelius Kelly for welcoming and hosting the Imperial-based research team in CIPPRes Clinic. We also thank all the volunteers for their participation.

## 1. Supplementary Data

The sections below give the full acquisition, preprocessing and analysis details for each modality, complementing the summary provided in the Methods of the main manuscript.

### 1.1. PET acquisition and analysis

We acquired dynamic PET data over a 90-minute scanning window. Continuous arterial blood sampling was performed via arterial cannulation to generate a whole blood activity curve. Discrete plasma samples were collected at defined intervals to estimate the plasma-over-blood ratio and to determine the parent fraction by high-performance liquid chromatography (HPLC), enabling construction of the metabolite-corrected arterial input function. All image data were analysed using the Perceptive Discovery in-house PET quantification tool, MIAKAT, implemented in MATLAB using SPM12 for image registration and segmentation. Dynamic PET images were registered to the participant’s T1-weighted MRI and corrected for motion. A 1-tissue compartment model for reversible binding was applied to estimate the total volume of distribution (V_T_) and registered to MNI152 space. V_T_ values were extracted for the regions of the Schaefer atlas (100 cortical-only parcels) (*33*) and of the CIC atlas (in-house atlas developed by Perceptive Discovery), which consists of a total of 121 grey matter regions, both cortical and subcortical (*34*). To limit partial-volume effects, regional V_T_ was averaged over grey-matter voxels only, defined as voxels with a grey-matter probability > 0.2 in each participant’s FSL FAST segmentation of the T1-weighted image, registered to MNI152 space. To obtain non-displaceable binding potential (BP_ND_), the V_T_ from each region was normalised by the V_T_ of the reference region, the centrum semiovale (CS), by computing the Distribution Volume Ratio-1 (DVR-1), obtained by dividing the region V_T_ by the CS V_T_ (reference region) minus one.

### 1.2. Resting-state fMRI acquisition and analysis

Resting-state fMRI data were acquired using an echo-planar imaging (EPI) sequence sensitive to blood-oxygen-level-dependent (BOLD) contrast with the following parameters: TR = 2,000 ms; TE = 30 ms; flip angle = 80°; in-plane resolution = 3 × 3 mm² (64 × 64 matrix); slice thickness = 3.6 mm; 36 axial slices; 240 volumes (10 minutes). A T1-weighted IR-SPGR structural image was also acquired (TI = 400 ms; TE minimum; flip angle = 11°; 256 × 256 matrix; 1 mm isotropic voxels; sagittal slices) to provide an anatomical reference.

MRI images were transformed from DICOM to NIfTI file format using dicom2nifti (*35*) and stored in BIDS format. fMRI preprocessing began with fMRIPrep (*36*), run via a Singularity image, which registered the T1w structural image to the BOLD image using FSL’s boundary-based registration (BBR) (*37*) with 12 degrees of freedom. Time series of 24 motion-related confounds (translation and rotation in the x, y, and z planes, their temporal and spatial derivatives, and framewise displacement (mm)) were computed using FSL (*70*) as part of the fMRIPrep pipeline and regressed out of the fMRI data using AFNI’s *3dTproject* function (*38*). 3dTproject uses linear regression to concurrently apply a band-pass filter (0.01–0.20 Hz), regress out the motion confounds and perform spatial smoothing with a Gaussian kernel of 8 mm full width at half maximum (FWHM).

After preprocessing, two atlases were transformed into subject space: the Schaefer 100-parcellation atlas and the CIC atlas. We used FreeSurfer’s *mri_surf2surf* function to perform surface-based registration and annotation mapping from the Schaefer atlas annotation in fsaverage space to each participant’s native surface space (*71*). This assigns a label to each vertex corresponding to the Schaefer region to which it belongs. Then, FreeSurfer’s *mri_aparc2aseg* function maps vertex labels from the cortical surface to the automatic segmentation volume. In the Schaefer atlas all region labels are also assigned a label corresponding to one of the seven resting state networks (*33*). Because no annotated, surface-based CIC atlas exists, we transformed the CIC atlas from standard MNI volumetric space into each participant’s native volumetric, functional space using a two-stage procedure. First, the participant’s skull-stripped T1-weighted brain image was nonlinearly registered to the MNI-space atlas template using Advanced Normalization Tools (ANTs 2.5.0) (*39*). Registration comprised an initial affine alignment followed by symmetric diffeomorphic normalisation (SyN) with a gradient step size of 0.5. Similarity was optimised with a Pearson correlation metric (weight = 1, radius = 2), with Gaussian regularisation of the update and total deformation fields (σ = 2 and 0.5 voxels, respectively). Optimisation ran over three resolution levels (100, 300 and 120 iterations, from coarse to fine), and image intensities were histogram-matched before registration. We applied the resulting affine and inverse warp fields to bring the atlas into the participant’s anatomical space, using nearest-neighbour interpolation to preserve discrete region labels. Finally, the atlas was resampled onto the grid of the participant’s resting-state BOLD reference image using FSL FLIRT (*70*) and the sform/qform header information to map between the two images, again with nearest-neighbour interpolation.

Time series from both atlases were extracted using the fslmeants function, which computes the average time course across all voxels in a region. Pairwise functional connectivity was computed as Spearman’s rho between all pairs of time series. The functional weighted degree (FWD) of each region was defined as the row-wise average of its Spearman’s correlation connectivity matrix.

### 1.3. DWI acquisition and analysis

DWI was acquired using an axial spin-echo EPI NODDI sequence. Two diffusion-weighted shells were acquired in the posterior-to-anterior (PA) phase-encoding direction: one at b = 1,000 s/mm² and one at b = 2,000 s/mm², each with 60 diffusion directions and identical acquisition parameters (TR/TE = 4,900/92 ms; FOV = 21.0 cm; slice thickness = 2.5 mm; matrix = 108 × 108), along with six *b* = 0 images. Three additional *b* = 0 images with reversed phase encoding (anterior-to-posterior) were acquired for susceptibility distortion correction.

DWI data were pre-processed using a pipeline combining MRtrix3 (*40*), FSL (*41*), and ANTs (*42*). Preprocessing steps included: (1) denoising of each shell separately; (2) Gibbs ringing correction; (3) susceptibility-induced distortion correction using pairs of opposing phase-encoding b0 images with FSL TOPUP; (4) eddy current and head motion correction with FSL EDDY, producing eddy-corrected volumes and rotated gradient vectors; (5) bias field correction using ANTs N4BiasFieldCorrection applied to the EDDY output. Quality control was performed by inspecting the EDDY outlier metrics.

Standard DTI metrics, including fractional anisotropy (FA), mean diffusivity (MD), axial diffusivity (AD), and radial diffusivity (RD), were extracted by fitting the diffusion tensor to the b = 1,000 s/mm² shell using FSL DTIFIT. NODDI model fitting was performed in MATLAB using the NODDI toolbox (*67*) to derive the orientation dispersion index (ODI). To reduce partial-volume contamination by cerebrospinal fluid, regional NODDI metrics were computed as tissue-weighted means (*43*): within each region, ODI was weighted voxel-wise by the tissue fraction (1 − FISO, where FISO is the NODDI isotropic volume fraction), i.e. ODI_TW = Σ ODI·(1 − FISO) / Σ (1 − FISO). Tissue-weighted ODI was used in all analyses. Subject-level DTI and NODDI maps were registered to 2 mm isotropic MNI152 standard space using a two-step ANTs registration pipeline: a rigid-body transform from DWI to the participant’s T1-weighted image, followed by a nonlinear SyN diffeomorphic transform from the T1 to the MNI152 template. Mean ODI, MD and FA values were extracted for each participant for each region of the Schaefer atlas and the CIC atlas.

Whole-brain structural connectomes were reconstructed for each participant and session following the approach described by Koller et al. (2024) (*44*), implemented in MRtrix3. Tissue-specific fibre orientation distributions (FODs) were estimated using multi-shell multi-tissue constrained spherical deconvolution (msmt-CSD), with white matter, grey matter, and CSF response functions derived using the Dhollander algorithm. FODs were intensity-normalised across tissue compartments using mtnormalise to ensure inter-subject comparability. Anatomically constrained tractography (ACT) was employed, using a five-tissue-type (5TT) segmentation image derived from FreeSurfer’s recon-all output and coregistered to DWI native space. Streamlines were seeded exclusively from the grey matter–white matter interface. Whole-brain probabilistic tractography was performed using the iFOD2 algorithm, generating 10 million streamlines per session (FOD amplitude cutoff = 0.06; maximum streamline length = 250 mm; backtracking enabled). To correct for inherent tractography biases that over-represent short, superficial fibres, streamline weights were estimated using SIFT2 (*45*). Tractography was performed entirely in native diffusion space to avoid the spatial distortion associated with warping tractograms; the Schaefer and CIC atlases, both defined in MNI152 space, were instead inverse-warped into each participant’s native diffusion space using the concatenated inverse of the rigid diffusion-to-T1 and nonlinear T1-to-MNI152 (ANTs SyN) transforms, with nearest-neighbour interpolation to preserve discrete ROI labels. A symmetric, SIFT2-weighted structural connectivity matrix was then computed for each participant using tck2connectome, with each matrix entry Wij representing the sum of SIFT2 weights across all streamlines connecting native-space voxels corresponding to atlas regions i and j, providing an index of fibre density between region pairs. The structural weighted degree (SWD) of each region was defined as the row-wise sum of its SIFT2-weighted connectivity matrix; for the CIC atlas, the sum was restricted to connections with the 121 grey-matter regions (excluding white-matter and ventricular labels).

### 1.4. EEG acquisition and analysis

#### 1.4.1. Roving mismatch negativity (MMN) task

The roving MMN paradigm followed the protocol described by Sumner et al. (2020) (*46*). Participants passively listened to sequences of pure sine-wave tones delivered binaurally via headphones for 15 minutes, during the inter-block rest between the early and late post-tetanic vLTP blocks (see below).

The auditory and visual stimuli were administered using the Psychtoolbox (*47*) on MATLAB (2023a). A roving-standard design was employed, spanning 500–800 Hz in 50 Hz steps (500, 550, 600, 650, 700, 750, and 800 Hz). Each tone was a pure sine wave of 70 ms duration with 10 ms cosine rise/fall ramps to prevent auditory clicks. Tones were presented in trains of 6-10 stimuli (train length = 5 + a random draw of 1-5 additional standards), giving a total of 500 trains across the session. The first tone in each new train constituted the deviant (tone position S1), and the sixth tone constituted the standard (tone position S6). The frequency for each new train was drawn randomly from the seven available frequencies, constrained to differ from that of the immediately preceding train. The stimulus onset asynchrony (SOA) between consecutive tones was approximately 500 ms (yielding a silent gap of ∼430 ms following the 70 ms tone).

To maintain alertness without directing attention to the tones, participants simultaneously performed a visual fixation distractor task: a central fixation cross underwent brief pseudorandomised luminance changes (every 30-60 seconds), and participants were instructed to press a button when they detected a change. Audio volume was set to a comfortable level for each participant at the start of the task and recorded to ensure consistency across sessions.

The MMN data analyses were implemented in MATLAB (2023a) using the FieldTrip toolbox (*48*). Raw continuous MMN data were bandpass-filtered (0.1-30 Hz; 2nd-order Butterworth high-pass, 30 Hz low-pass). Epochs were defined for each of the six tone positions (S1 to S6) with a pre-stimulus window of −100 ms and a post-stimulus window of +400 ms. Baseline correction was applied over the pre-stimulus interval (−100 to 0 ms). Artefact rejection followed the same protocol as for the vLTP data (described below).

The choice of electrodes and time windows of interest for the ERPs was based on previous literature (*49*). The *a priori* electrode of interest was Fz. The MMN amplitude was quantified as the mean amplitude (µV) within a 100–300 ms post-stimulus window. Component windows and electrode selections were confirmed by visual inspection of the grand-average MMN difference waveform across all participants. MMN repetition suppression was quantified as the linear slope of amplitude change across the six tone positions (S1 to S6) using first-order polynomial fitting, providing a parametric index of the rate of adaptation from deviant to standard.

#### 1.4.2. Visual long-term potentiation (vLTP) task

The vLTP task followed the protocol described by Sumner et al. (2020) (*50*). The visual stimuli were administered using the Psychtoolbox (*47*) on MATLAB (2023a). Participants viewed circular sinusoidal grating stimuli (8 degrees of visual angle; spatial frequency 8 cycles per degree; ∼33 ms stimulus duration) presented on a mid-grey background, with a central red fixation dot maintained throughout. Two orthogonal orientations were used, vertical and horizontal, counterbalanced such that odd-numbered participants received horizontal tetanus and even-numbered participants received vertical tetanus. The inter-stimulus interval (ISI) was pseudorandomly jittered around the target frequency (3 steady : 1 short : 1 long ISI ratio) to minimise stimulus predictability.

The full vLTP session comprised the following sequential blocks:

(1) Pre-tetanus baseline (1 Hz stimulation, ∼4 minutes): 240 trials (120 vertical + 120 horizontal gratings, pseudorandomised), presented at 1 Hz with a jittered ISI of ∼967 ms (±33 ms). Both stimulus orientations were presented at equal frequency in this and all post-tetanic assessment blocks.
(2) Tetanic stimulation block (∼8.4 Hz, ∼2 minutes): 1,000 flashes of the tetanus-assigned grating orientation only (horizontal for odd-numbered participants, vertical for even-numbered participants), delivered at a nominal target frequency of 9 Hz, intended to induce LTP-like synaptic potentiation. Because stimulus onsets were locked to the 60 Hz monitor refresh (16.7 ms per refresh cycle), the 111 ms target period (6.67 refresh cycles) could not be reproduced exactly, and the achieved presentation rate was ∼8.4 Hz (ISI ∼86 ms, jittered by ±1 refresh cycle, ∼16 ms).
(3) 2-minute eyes-closed rest break.
(4) Early post-tetanic assessment block (1 Hz, ∼4 minutes): 240 trials repeating the pre-tetanus baseline structure, recorded approximately 2 minutes after the end of the tetanic block.
(5) Inter-block rest (after the early post-tetanus block), during which the roving MMN task was administered (see above).
(6) Late post-tetanic assessment block (1 Hz, ∼4 minutes): 240 trials repeating the pre-tetanus baseline structure, recorded approximately 30 minutes after the tetanic stimulation block.

The vLTP data analyses were implemented in MATLAB (2023a) using the FieldTrip toolbox (*48*). Raw continuous data were bandpass-filtered between 0.1 Hz and 30 Hz. Epochs were defined relative to stimulus onset triggers, with a pre-stimulus window of −200 ms and a post-stimulus window of +500 ms. Baseline correction was applied over the pre-stimulus interval (−200 to 0 ms). Artefact rejection proceeded in two stages. First, blinks and ocular artefacts which overlapped with the onset of stimulus presentation were identified semi-automatically using a Z-value threshold applied to frontal electrodes (Fp1, Fp2; band-pass 0.3–1 Hz, Hilbert envelope), followed by manual verification and rejection. Further noisy epochs were manually identified and rejected. Second, bad channels were identified and excluded by visual inspection using a graphical channel selection interface, after which rejected channels were interpolated using spherical spline interpolation. Then, independent component analysis (ICA) was performed using the ‘runica’ algorithm to reject residual ocular, heartbeat, and other non-brain components. The data were then re-referenced to the common average.

Event-related potentials (ERPs) were computed for each block condition (Pre-tetanus, Early Post-tetanus, Late Post-tetanus) by averaging all accepted epochs from the tetanised stimulus only. The primary ERP outcome measures were the N1b component for early post-tetanic potentiation and the P2 component for late-phase potentiation. The choice of electrodes and time windows of interest for the ERPs was based on previous literature (*51*, *52*). For N1b, the *a priori* electrodes of interest were PO3 and PO4. For these electrodes, the grand-average of the N1b potentiation was visually inspected, and the mean amplitude (µV) was extracted from the 80–150 ms time window. For the P2, the *a priori* electrode of interest was POz. The grand-average of the P2 potentiation was visually inspected, and the peak amplitude (µV) was extracted within the 150–300 ms time window. Late-phase P2 potentiation (ΔP2-Late) was computed for each participant as the difference in P2 peak amplitude between the late post-tetanic and the pre-tetanus blocks.

### 1.5. Regions of interest for the EEG-PET analyses

Regions of interest for the EEG-PET analyses were defined from the Schaefer and CIC atlases a priori, based on the known anatomical generators of each paradigm (*49*, *51*). The frontotemporal group (MMN) comprised 22 CIC regions (left and right anterior and posterior superior temporal gyrus, frontal operculum, anterior and posterior dorsolateral frontal cortex, anterior and posterior medial frontal cortex, hippocampus and amygdala, and the midline subcallosal, anterior, dorsal anterior and posterior cingulate gyri) and the 43 Schaefer parcels belonging to the following sub-networks: Control (lateral, medial-posterior and ventral prefrontal, orbitofrontal, cingulate and temporal), Salience/Ventral Attention (lateral prefrontal, medial, and frontal operculum/insula), Default (prefrontal, dorsal and medial prefrontal, ventral prefrontal, precuneus/posterior cingulate, temporal and parahippocampal) and Limbic (orbitofrontal and temporal pole). The occipital group (vLTP) comprised 12 CIC regions (left and right calcarine cortex, cuneus, lingual gyrus, occipital pole, occipital fusiform gyrus and temporo-occipital fusiform cortex) and the 17 parcels of the Schaefer visual network.

For each group, a volume correction was applied: V_T_ was averaged across the constituent regions weighted by their number of grey-matter voxels, and the group BP_ND_ was computed from this volume-weighted V_T_ (BP_ND_ = V_T_/V_T,CS_ − 1), so that large and small regions contribute in proportion to their size.

### 1.6. Statistical models

All statistical models were fitted in R with the lme4, lmerTest and emmeans packages (Kenward-Roger degrees of freedom computed with pbkrtest); intraclass correlation coefficients were computed with the psych package. Region-level analyses used one value per subject, scan and region (Schaefer: 100 parcels; CIC: 121 grey-matter regions).

Drug effects were tested on subject-level means: each region-level measure was first averaged across regions (separately for cortical and subcortical CIC regions), and the subject means were modelled as Outcome ∼ Time + (1 | Subject_ID), fitted by maximum likelihood. For the CIC atlas, the cortical and subcortical p-values were FDR-corrected.

For the region-level PET-MRI associations, each predictor X was decomposed, within each atlas and scan, into three components: X_subject_, the mean of X across regions for each subject; X_region_, the mean of X across subjects for each region; and X_within_ = X - X_subject_ - X_region_ + the grand mean of X. All three components were centred and divided by the pooled standard deviation of X, so that their coefficients are on a common scale (change in the outcome per pooled standard deviation of the predictor). Global models, Outcome ∼ X_within_ + X_subject_ + X_region_ + (1 | Subject_ID) + (1 | ROI_Label), were fitted by maximum likelihood. Region-stratified models added the interaction between X_within_ and the atlas subdivision (the seven Schaefer networks, or cortical vs subcortical CIC regions) and were fitted by restricted maximum likelihood; subdivision-specific slopes were estimated as marginal trends with Kenward-Roger degrees of freedom and FDR-corrected across the subdivisions of each model, and the overall difference between subdivisions was tested with the F-test of the interaction. The same model, with one MRI measure as the outcome and another as the predictor, was used to assess collinearity between MRI measures. The stability of the associations between scans was tested by fitting both scans together, with X decomposed within each scan: Outcome ∼ (X_within_ + X_subject_ + X_region_) × Scan + (1 | Subject_ID/Scan) + (1 | ROI_Label).

Test-retest reliability was quantified with ICC(A,1) (two-way model, absolute agreement, single measure) on one value per subject and scan. The change in each subject-level EEG-PET correlation between scans was assessed with Zou’s 95% confidence interval for the difference between two dependent, non-overlapping correlations (*54*).

## 2. Supplementary Results

### 2.1. Effects of ketamine and DMT on PET, MRI and EEG measures

To test whether each measure changed after a single dose of ketamine or DMT, the post-drug re-test scan was compared with the baseline scan in the whole sample (n = 18; ketamine and DMT combined) with linear mixed models fitted to subject-level means (one value per subject and scan, averaged across regions): Outcome ∼ Time + (1 | Subject_ID). Results are reported for the Schaefer atlas (primary analysis, as in the main manuscript), for the CIC atlas (overall, and separately for cortical and subcortical regions, FDR-corrected across the two region types), and for the subject-level EEG markers (Supplementary Table 1).

Schaefer atlas. No significant changes were observed in BP_ND_ (β = -0.030, p = 0.615), V_T_ (β = 0.955, p = 0.120), ODI (β = 0.0007, p = 0.443), SWD (β = -94.6, p = 0.771) or FWD (β = -0.017, p = 0.574).

CIC atlas. BP_ND_ did not change significantly overall (β = -0.003, p = 0.961), nor within cortical (β = -0.007, p_FDR_ = 0.954) or subcortical (β = 0.003, p_FDR_ = 0.954) regions. V_T_ showed a trend-level increase overall (β = 0.941, p = 0.081), in both cortical (β = 0.996, p_FDR_ = 0.084) and subcortical (β = 0.857, p_FDR_ = 0.084) regions. ODI did not change significantly overall (β = 0.001, p = 0.566), nor within cortical (β = 0.001, p_FDR_ = 0.759) or subcortical (β = 0.001, p_FDR_ = 0.759) regions. This dissociation between a trend-level V_T_ increase and a null BP_ND_ finding (as reported in the main manuscript) is consistent with a discrepancy between these two metrics that we have previously reported in the ketamine cohort (*72*). Here, in the cohort enlarged with the DMT participants, V_T_ also showed a trend-level increase in the reference region (centrum semiovale). Because BP_ND_ is computed relative to this reference region, a non-specific, drug-related rise in V_T_ that is not confined to grey matter would be expected to attenuate genuine grey-matter increases in BP_ND_. As BP_ND_ corrects for non-displaceable binding and is therefore considered a more specific index of synaptic density than V_T_, and since the primary aim of the current analysis is to characterise cross-modal relationships between synaptic density and structural/functional plasticity markers, rather than to re-establish the overall effect of drug administration on any single metric, BP_ND_ was used as the primary synaptic density outcome throughout the current analysis. SWD did not change significantly overall (β = -86.7, p = 0.823), nor within cortical (p_FDR_ = 0.763) or subcortical (p_FDR_ = 0.763) regions. MD (p = 0.521) and FA (p = 0.215) did not change significantly overall, nor within cortical or subcortical regions (all p_FDR_ > 0.200). FWD did not change significantly overall (β = -0.023, p = 0.362), nor within cortical (p_FDR_ = 0.482) or subcortical (p_FDR_ = 0.425) regions.

EEG. Neither the MMN repetition suppression slope (β = 0.019, p = 0.229) nor late-phase P2 potentiation (ΔP2-Late; β = -1.736, p = 0.100) changed significantly between the baseline and re-test scans.

**Supplementary Table 1.** Change in each measure between the baseline and the post-drug re-test scan (n = 18; ketamine, n = 11; DMT, n = 7). Linear mixed models fitted by maximum likelihood to subject-level means (one value per subject and scan, averaged across regions): Outcome ∼ Time + (1 | Subject_ID). β is the re-test minus baseline difference, in the raw units of each measure. For the CIC atlas, cortical and subcortical p-values are FDR-corrected across the two region types. RS, repetition suppression; ΔP2-Late, late-phase P2 potentiation.

| Atlas | Measure | Regions | $\beta$ | SE | p |
| --- | --- | --- | --- | --- | --- |
| Schaefer | BP<sub>ND</sub> | All regions | -0.0301 | 0.0587 | 0.615 |
| Schaefer | V<sub>T</sub> | All regions | 0.955 | 0.584 | 0.120 |
| Schaefer | ODI | All regions | $6.72 \times 10^{-4}$ | $8.56 \times 10^{-4}$ | 0.443 |
| Schaefer | SWD | All regions | -94.6 | 319.5 | 0.771 |
| Schaefer | FWD | All regions | -0.0169 | 0.0295 | 0.574 |
| CIC | BP<sub>ND</sub> | All regions | -0.00263 | 0.0535 | 0.961 |
| CIC | BP<sub>ND</sub> | Cortical | -0.00682 | 0.0538 | 0.954 |
| CIC | BP<sub>ND</sub> | Subcortical | 0.00318 | 0.0543 | 0.954 |
| CIC | V<sub>T</sub> | All regions | 0.941 | 0.508 | 0.081 |
| CIC | V<sub>T</sub> | Cortical | 0.996 | 0.535 | 0.084 |
| CIC | V<sub>T</sub> | Subcortical | 0.857 | 0.468 | 0.084 |
| CIC | ODI | All regions | $6.01 \times 10^{-4}$ | 0.00103 | 0.566 |
| CIC | ODI | Cortical | $6.50 \times 10^{-4}$ | $9.13 \times 10^{-4}$ | 0.759 |
| CIC | ODI | Subcortical | $5.28 \times 10^{-4}$ | 0.00170 | 0.759 |
| CIC | MD | All regions | $1.89 \times 10^{-6}$ | $2.89 \times 10^{-6}$ | 0.521 |
| CIC | MD | Cortical | $5.08 \times 10^{-6}$ | $3.72 \times 10^{-6}$ | 0.376 |
| CIC | MD | Subcortical | $-2.80 \times 10^{-6}$ | $4.32 \times 10^{-6}$ | 0.525 |
| CIC | FA | All regions | -0.00153 | 0.00119 | 0.215 |
| CIC | FA | Cortical | -0.00169 | $9.83 \times 10^{-4}$ | 0.204 |
| CIC | FA | Subcortical | -0.00128 | 0.00180 | 0.488 |
| CIC | SWD | All regions | -86.7 | 382.2 | 0.823 |
| CIC | SWD | Cortical | -234.4 | 444.1 | 0.763 |
| CIC | SWD | Subcortical | 130.3 | 426.3 | 0.763 |
| CIC | FWD | All regions | -0.0232 | 0.0248 | 0.362 |
| CIC | FWD | Cortical | -0.0188 | 0.0261 | 0.482 |
| CIC | FWD | Subcortical | -0.0299 | 0.0235 | 0.425 |
| EEG | MMN RS slope | Fz | 0.0189 | 0.0152 | 0.229 |
| EEG | ΔP2-Late | POz | -1.74 | 1.00 | 0.100 |

### 2.2. Collinearity between MRI and EEG measures

To establish whether ODI, SWD and FWD are independent predictors of BP_ND_, collinearity between the MRI measures was assessed with the same within-between model, with one MRI measure as the outcome and the other as the predictor; the within-region term is reported (Supplementary Table 2). At baseline, the individual MMN repetition suppression slope and late-phase P2 potentiation (ΔP2-Late) were not correlated (Pearson’s r = 0.13, p = 0.617, n = 18), indicating that the two EEG markers are statistically independent.

**Supplementary Table 2.**
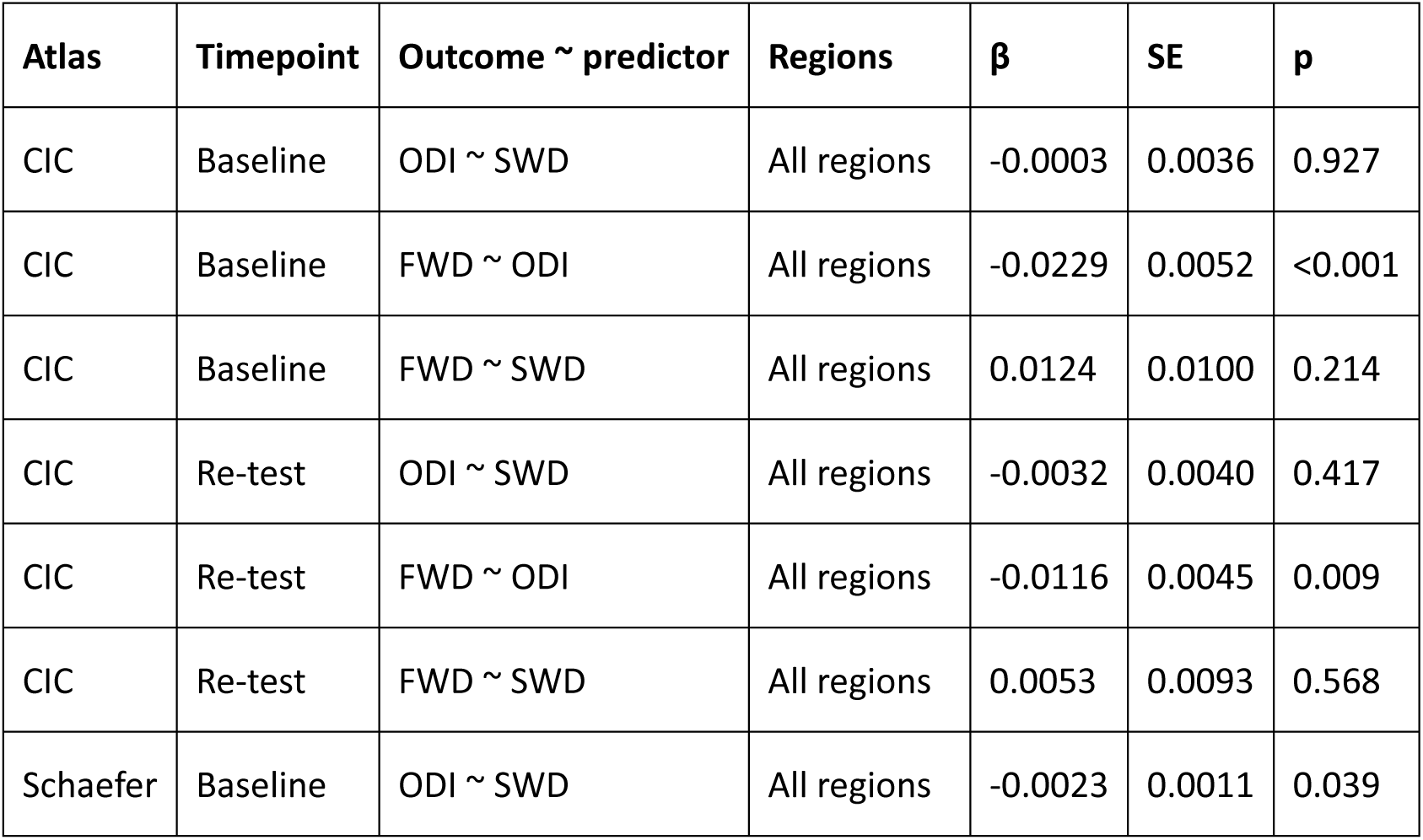
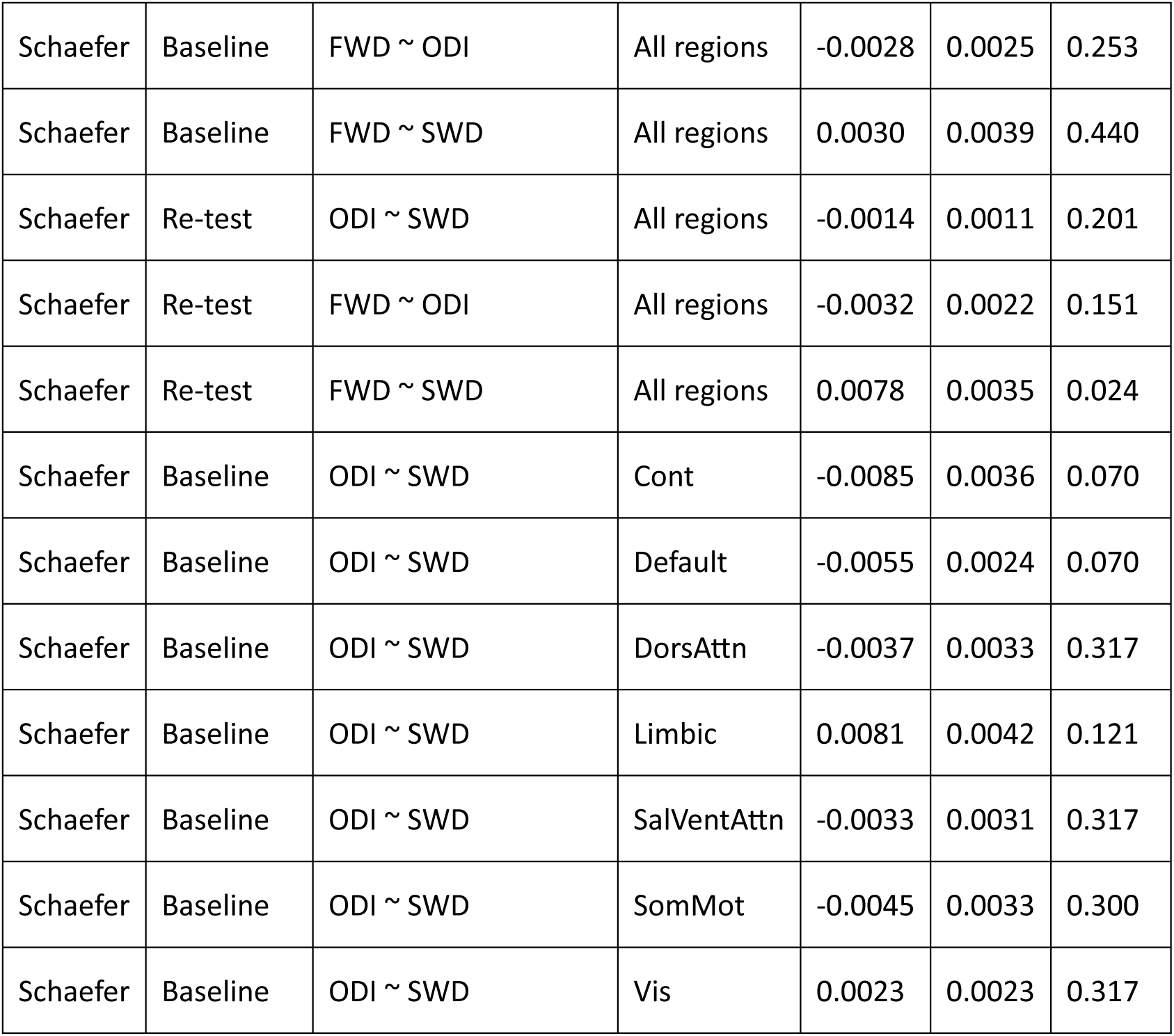
Collinearity between MRI measures. Within-between linear mixed models with one MRI measure as the outcome and the other as the predictor: Outcome ∼ X_within_ + X_subject_ + X_region_ + (1 | Subject_ID) + (1 | ROI_Label). The within-region term (X_within_; change in the outcome per pooled SD of the predictor) is reported. Network-specific slopes (Schaefer atlas, baseline) were estimated from the region-stratified model and FDR-corrected across the seven networks. Networks: Cont, control; Default, default mode; DorsAttn, dorsal attention; SalVentAttn, salience/ventral attention; SomMot, somatomotor; Vis, visual.

| Atlas | Timepoint | Outcome ∼ predictor | Regions | $\beta$ | SE | p |
| --- | --- | --- | --- | --- | --- | --- |
| CIC | Baseline | ODI ∼ SWD | All regions | -0.0003 | 0.0036 | 0.927 |
| CIC | Baseline | FWD ∼ ODI | All regions | -0.0229 | 0.0052 | <0.001 |
| CIC | Baseline | FWD ∼ SWD | All regions | 0.0124 | 0.0100 | 0.214 |
| CIC | Re-test | ODI ∼ SWD | All regions | -0.0032 | 0.0040 | 0.417 |
| CIC | Re-test | FWD ∼ ODI | All regions | -0.0116 | 0.0045 | 0.009 |
| CIC | Re-test | FWD ∼ SWD | All regions | 0.0053 | 0.0093 | 0.568 |
| Schaefer | Baseline | ODI ∼ SWD | All regions | -0.0023 | 0.0011 | 0.039 |
| Schaefer | Baseline | FWD ~ ODI | All regions | -0.0028 | 0.0025 | 0.253 |
| Schaefer | Baseline | FWD ~ SWD | All regions | 0.0030 | 0.0039 | 0.440 |
| Schaefer | Re-test | ODI ~ SWD | All regions | -0.0014 | 0.0011 | 0.201 |
| Schaefer | Re-test | FWD ~ ODI | All regions | -0.0032 | 0.0022 | 0.151 |
| Schaefer | Re-test | FWD ~ SWD | All regions | 0.0078 | 0.0035 | 0.024 |
| Schaefer | Baseline | ODI ~ SWD | Cont | -0.0085 | 0.0036 | 0.070 |
| Schaefer | Baseline | ODI ~ SWD | Default | -0.0055 | 0.0024 | 0.070 |
| Schaefer | Baseline | ODI ~ SWD | DorsAttn | -0.0037 | 0.0033 | 0.317 |
| Schaefer | Baseline | ODI ~ SWD | Limbic | 0.0081 | 0.0042 | 0.121 |
| Schaefer | Baseline | ODI ~ SWD | SalVentAttn | -0.0033 | 0.0031 | 0.317 |
| Schaefer | Baseline | ODI ~ SWD | SomMot | -0.0045 | 0.0033 | 0.300 |
| Schaefer | Baseline | ODI ~ SWD | Vis | 0.0023 | 0.0023 | 0.317 |

### 2.3. Overview of the multimodal association analyses

The following sections report all associations between synaptic terminal SV2A density ([¹¹C]-UCBJ BP_ND_, and V_T_ as a sensitivity outcome) and the MRI (ODI, MD, FA, SWD, FWD) and EEG (MMN repetition suppression, late-phase P2 potentiation) measures. Each result is reported separately for the baseline scan and the post-drug re-test scan, and for the Schaefer atlas (100 cortical parcels, seven networks; primary) and the CIC atlas (121 cortical and subcortical grey-matter regions; validation). Region-level MRI associations were tested with the within-between linear mixed models described in the Supplementary Methods: the global model gives the within-region, between-subject and between-region terms, and the region-stratified model gives the network-specific (Schaefer) or cortical/subcortical (CIC) within-region slopes. Subject-level EEG associations were tested with Pearson correlations (n = 18) and, at baseline, with the EEG marker added as a covariate to the ODI-BP_ND_ model.

#### 2.3.1. Relationship between synaptic terminal SV2A density and neurite complexity

Linear mixed models were fitted with synaptic density (BP_ND_ or V_T_) as the outcome and ODI as the predictor. ODI was split into a within-component (ODI minus the subject mean and the region mean, plus the grand mean) and the subject and region means, all scaled by the pooled SD of ODI, and all three were entered as fixed effects with random intercepts for Subject and ROI: PET outcome (BP_ND_/V_T_) ∼ ODI_within_ + ODI_subject_ + ODI_region_ + (1 | Subject_ID) + (1 | ROI_Label). The ODI_within_ coefficient, reported throughout, estimates the association across individuals within regions, free of between-subject and between-region associations; the ODI_region_ and ODI_subject_ coefficients reflect the between-region (spatial) and between-subject associations, respectively. A region-stratified model was additionally fitted to test whether the ODI-synaptic density relationship differed between cortical and subcortical areas (CIC atlas) or across the seven functional networks (Schaefer atlas): PET outcome (BP_ND_/V_T_) ∼ ODI_within_ * Network + ODI_subject_ + ODI_region_ + (1 | Subject_ID) + (1 | ROI_Label). Region-specific slopes of ODI_within_ were extracted with emtrends (Kenward-Roger degrees of freedom) and p-values corrected using the false discovery rate (FDR). Results are reported in Supplementary Table 3.

**Supplementary Table 3.** Associations between neurite orientation dispersion index (ODI) and synaptic terminal SV2A density (BP_ND_ and V_T_) in the CIC atlas (cortical and subcortical regions) and the Schaefer atlas (100 parcels, seven networks), at baseline and re-test. Within-between linear mixed models with random intercepts for subject and region. Within-region (global): within-region term of the global model (maximum likelihood); Between-subject and Between-region: subject-mean and region-mean terms of the same model; Within-region (stratified): cortical/subcortical or network-specific within-region slopes from the region-stratified model (restricted maximum likelihood; Kenward-Roger degrees of freedom), with p-values FDR-corrected across the subdivisions of each model. β is the change in BP_ND_ or V_T_ per pooled SD of ODI; because subject means vary much less than individual values, between-subject β values are correspondingly large. Networks: Cont, control; Default, default mode; DorsAttn, dorsal attention; SalVentAttn, salience/ventral attention; SomMot, somatomotor; Vis, visual.

| Atlas | Timepoint | Outcome | Term | Region | $\beta$ | SE | p |
| --- | --- | --- | --- | --- | --- | --- | --- |
| CIC | Baseline | $BP_{ND}$ | Within-region (global) | - | 0.139 | 0.020 | <0.001 |
|  |  |  | Between-subject | - | 0.950 | 0.902 | 0.306 |
|  |  |  | Between-region | - | 0.866 | 0.056 | <0.001 |
|  |  |  | Within-region (stratified) | Cortical | 0.085 | 0.036 | 0.020 |
|  |  |  | Within-region (stratified) | Subcortical | 0.166 | 0.025 | <0.001 |
| CIC | Baseline | $V_T$ | Within-region (global) | - | 0.517 | 0.103 | <0.001 |
|  |  |  | Between-subject | - | 20.350 | 9.263 | 0.041 |
|  |  |  | Between-region | - | 3.571 | 0.233 | <0.001 |
|  |  |  | Within-region (stratified) | Cortical | 0.099 | 0.184 | 0.590 |
|  |  |  | Within-region (stratified) | Subcortical | 0.721 | 0.127 | <0.001 |
| CIC | Re-test | $BP_{ND}$ | Within-region (global) | - | 0.136 | 0.020 | <0.001 |
|  |  |  | Between-subject | - | 0.102 | 0.815 | 0.902 |
|  |  |  | Between-region | - | 0.853 | 0.057 | <0.001 |
|  |  |  | Within-region (stratified) | Cortical | 0.176 | 0.037 | <0.001 |
|  |  |  | Within-region (stratified) | Subcortical | 0.118 | 0.024 | <0.001 |
| CIC | Re-test | V<sub>T</sub> | Within-region (global) | - | 0.657 | 0.104 | <0.001 |
|  |  |  | Between-subject | - | 14.441 | 9.476 | 0.145 |
|  |  |  | Between-region | - | 3.709 | 0.249 | <0.001 |
|  |  |  | Within-region (stratified) | Cortical | 0.843 | 0.193 | <0.001 |
|  |  |  | Within-region (stratified) | Subcortical | 0.574 | 0.127 | <0.001 |
| Schaefer | Baseline | BP<sub>ND</sub> | Within-region (global) | - | 0.036 | 0.007 | <0.001 |
|  |  |  | Between-subject | - | 0.270 | 0.376 | 0.482 |
|  |  |  | Between-region | - | 0.006 | 0.064 | 0.926 |
|  |  |  | Within-region (stratified) | Cont | 0.047 | 0.020 | 0.037 |
|  |  |  | Within-region (stratified) | Default | 0.004 | 0.017 | 0.902 |
|  |  |  | Within-region (stratified) | DorsAttn | 0.067 | 0.019 | 0.004 |
|  |  |  | Within-region (stratified) | Limbic | 0.075 | 0.031 | 0.035 |
|  |  |  | Within-region (stratified) | SalVentAttn | -0.003 | 0.025 | 0.902 |
|  |  |  | Within-region (stratified) | SomMot | 0.035 | 0.019 | 0.096 |
|  |  |  | Within-region (stratified) | Vis | 0.044 | 0.017 | 0.032 |
| Schaefer | Baseline | V<sub>T</sub> | Within-region (global) | - | 0.174 | 0.036 | <0.001 |
|  |  |  | Between-subject | - | 9.675 | 3.595 | 0.015 |
|  |  |  | Between-region | - | 0.032 | 0.264 | 0.904 |
|  |  |  | Within-region (stratified) | Cont | 0.277 | 0.096 | 0.014 |
|  |  |  | Within-region (stratified) | Default | 0.052 | 0.079 | 0.592 |
|  |  |  | Within-region (stratified) | DorsAttn | 0.269 | 0.092 | 0.014 |
|  |  |  | Within-region (stratified) | Limbic | 0.381 | 0.147 | 0.022 |
|  |  |  | Within-region (stratified) | SalVentAttn | 0.011 | 0.120 | 0.926 |
|  |  |  | Within-region (stratified) | SomMot | 0.168 | 0.092 | 0.095 |
|  |  |  | Within-region (stratified) | Vis | 0.170 | 0.080 | 0.058 |
| Schaefer | Re-test | BP<sub>ND</sub> | Within-region (global) | - | 0.045 | 0.008 | <0.001 |
|  |  |  | Between-subject | - | 0.184 | 0.275 | 0.513 |
|  |  |  | Between-region | - | 0.009 | 0.066 | 0.894 |
|  |  |  | Within-region (stratified) | Cont | 0.050 | 0.020 | 0.023 |
|  |  |  | Within-region (stratified) | Default | 0.023 | 0.019 | 0.298 |
|  |  |  | Within-region (stratified) | DorsAttn | 0.054 | 0.019 | 0.017 |
|  |  |  | Within-region (stratified) | Limbic | 0.030 | 0.029 | 0.363 |
|  |  |  | Within-region (stratified) | SalVentAttn | 0.011 | 0.028 | 0.689 |
|  |  |  | Within-region (stratified) | SomMot | 0.052 | 0.019 | 0.017 |
|  |  |  | Within-region (stratified) | Vis | 0.068 | 0.019 | 0.002 |
| Schaefer | Re-test | $V_T$ | Within-region (global) | - | 0.178 | 0.037 | <0.001 |
|  |  |  | Between-subject | - | 9.302 | 2.737 | 0.003 |
|  |  |  | Between-region | - | 0.047 | 0.285 | 0.869 |
|  |  |  | Within-region (stratified) | Cont | 0.284 | 0.095 | 0.019 |
|  |  |  | Within-region (stratified) | Default | 0.102 | 0.088 | 0.344 |
|  |  |  | Within-region (stratified) | DorsAttn | 0.235 | 0.092 | 0.025 |
|  |  |  | Within-region (stratified) | Limbic | 0.063 | 0.138 | 0.757 |
|  |  |  | Within-region (stratified) | SalVentAttn | -0.030 | 0.130 | 0.821 |
|  |  |  | Within-region (stratified) | SomMot | 0.200 | 0.092 | 0.053 |
|  |  |  | Within-region (stratified) | Vis | 0.232 | 0.089 | 0.025 |

#### 2.3.2. Relationship between synaptic terminal SV2A density and mean diffusivity (MD)

Mean diffusivity (MD) was used as a marker of tissue microstructural integrity, with higher MD reflecting less-restricted diffusion consistent with reduced cellular density or tissue degradation. Linear mixed models were fitted with synaptic density (BP_ND_ or V_T_) as the outcome and MD as predictor, using the same within-between model as for ODI (predictor split into a within-component and subject and region means, with random intercepts for subject and ROI). Results are reported in Supplementary Table 4.

**Supplementary Table 4.** Associations between mean diffusivity (MD) and synaptic terminal SV2A density (BP_ND_ and V_T_) in the CIC atlas (cortical and subcortical regions) and the Schaefer atlas (100 parcels, seven networks), at baseline and re-test. Within-between linear mixed models with random intercepts for subject and region. Within-region (global): within-region term of the global model (maximum likelihood); Between-subject and Between-region: subject-mean and region-mean terms of the same model; Within-region (stratified): cortical/subcortical or network-specific within-region slopes from the region-stratified model (restricted maximum likelihood; Kenward-Roger degrees of freedom), with p-values FDR-corrected across the subdivisions of each model. β is the change in BP_ND_ or V_T_ per pooled SD of MD; because subject means vary much less than individual values, between-subject β values are correspondingly large. Networks: Cont, control; Default, default mode; DorsAttn, dorsal attention; SalVentAttn, salience/ventral attention; SomMot, somatomotor; Vis, visual.

| Atlas | Timepoint | Outcome | Term | Region | $\beta$ | SE | p |
| --- | --- | --- | --- | --- | --- | --- | --- |
| CIC | Baseline | BP<sub>ND</sub> | Within-region (global) | - | -<br>0.067 | 0.016 | <0.001 |
|  |  |  | Between-subject | - | -<br>1.316 | 0.551 | 0.028 |
|  |  |  | Between-region | - | -<br>0.551 | 0.087 | <0.001 |
|  |  |  | Within-region (stratified) | Cortical | 0.017 | 0.026 | 0.513 |
|  |  |  | Within-region (stratified) | Subcortical | -<br>0.117 | 0.020 | <0.001 |
| CIC | Baseline | V<sub>T</sub> | Within-region (global) | - | -<br>0.206 | 0.078 | 0.009 |
|  |  |  | Between-subject | - | -<br>6.548 | 6.920 | 0.357 |
|  |  |  | Between-region | - | -<br>2.267 | 0.361 | <0.001 |
|  |  |  | Within-region (stratified) | Cortical | 0.222 | 0.129 | 0.086 |
|  |  |  | Within-region (stratified) | Subcortical | -<br>0.461 | 0.099 | <0.001 |
| CIC | Re-test | BP<sub>ND</sub> | Within-region (global) | - | -<br>0.103 | 0.016 | <0.001 |
|  |  |  | Between-subject | - | -<br>0.709 | 0.465 | 0.145 |
|  |  |  | Between-region | - | -<br>0.535 | 0.088 | <0.001 |
|  |  |  | Within-region (stratified) | Cortical | -<br>0.020 | 0.024 | 0.413 |
|  |  |  | Within-region (stratified) | Subcortical | -<br>0.169 | 0.021 | <0.001 |
| CIC | Re-test | V<sub>T</sub> | Within-region (global) | - | -<br>0.647 | 0.081 | <0.001 |
|  |  |  | Between-subject | - | - 0.431 | 6.088 | 0.944 |
|  |  |  | Between-region | - | - 2.319 | 0.386 | <0.001 |
|  |  |  | Within-region (stratified) | Cortical | - 0.416 | 0.124 | <0.001 |
|  |  |  | Within-region (stratified) | Subcortical | - 0.828 | 0.109 | <0.001 |
| Schaefer | Baseline | BP<sub>ND</sub> | Within-region (global) | - | - 0.059 | 0.010 | <0.001 |
|  |  |  | Between-subject | - | - 0.296 | 0.233 | 0.219 |
|  |  |  | Between-region | - | - 0.361 | 0.048 | <0.001 |
|  |  |  | Within-region (stratified) | Cont | - 0.015 | 0.025 | 0.545 |
|  |  |  | Within-region (stratified) | Default | - 0.053 | 0.023 | 0.042 |
|  |  |  | Within-region (stratified) | DorsAttn | - 0.043 | 0.022 | 0.067 |
|  |  |  | Within-region (stratified) | Limbic | - 0.047 | 0.051 | 0.424 |
|  |  |  | Within-region (stratified) | SalVentAttn | - 0.123 | 0.030 | <0.001 |
|  |  |  | Within-region (stratified) | SomMot | - 0.090 | 0.023 | <0.001 |
|  |  |  | Within-region (stratified) | Vis | - 0.051 | 0.026 | 0.067 |
| Schaefer | Baseline | V<sub>T</sub> | Within-region (global) | - | - 0.269 | 0.046 | <0.001 |
|  |  |  | Between-subject | - | 0.196 | 2.717 | 0.943 |
|  |  |  | Between-region | - | -<br>1.493 | 0.197 | <0.001 |
|  |  |  | Within-region<br>(stratified) | Cont | -<br>0.049 | 0.120 | 0.686 |
|  |  |  | Within-region<br>(stratified) | Default | -<br>0.256 | 0.107 | 0.040 |
|  |  |  | Within-region<br>(stratified) | DorsAttn | -<br>0.151 | 0.103 | 0.200 |
|  |  |  | Within-region<br>(stratified) | Limbic | -<br>0.228 | 0.244 | 0.410 |
|  |  |  | Within-region<br>(stratified) | SalVentAttn | -<br>0.522 | 0.144 | 0.001 |
|  |  |  | Within-region<br>(stratified) | SomMot | -<br>0.468 | 0.108 | <0.001 |
|  |  |  | Within-region<br>(stratified) | Vis | -<br>0.253 | 0.122 | 0.066 |
| Schaefer | Re-test | BP<sub>ND</sub> | Within-region (global) | - | -<br>0.052 | 0.010 | <0.001 |
|  |  |  | Between-subject | - | -<br>0.301 | 0.164 | 0.082 |
|  |  |  | Between-region | - | -<br>0.375 | 0.051 | <0.001 |
|  |  |  | Within-region<br>(stratified) | Cont | -<br>0.003 | 0.025 | 0.899 |
|  |  |  | Within-region<br>(stratified) | Default | -<br>0.062 | 0.024 | 0.025 |
|  |  |  | Within-region<br>(stratified) | DorsAttn | -<br>0.023 | 0.024 | 0.404 |
|  |  |  | Within-region<br>(stratified) | Limbic | -<br>0.092 | 0.047 | 0.087 |
|  |  |  | Within-region<br>(stratified) | SalVentAttn | -<br>0.105 | 0.032 | 0.003 |
|  |  |  | Within-region (stratified) | SomMot | - 0.082 | 0.023 | 0.003 |
|  |  |  | Within-region (stratified) | Vis | - 0.037 | 0.027 | 0.232 |
| Schaefer | Re-test | V<sub>T</sub> | Within-region (global) | - | - 0.236 | 0.047 | <0.001 |
|  |  |  | Between-subject | - | - 1.476 | 2.224 | 0.515 |
|  |  |  | Between-region | - | - 1.623 | 0.221 | <0.001 |
|  |  |  | Within-region (stratified) | Cont | 0.060 | 0.120 | 0.720 |
|  |  |  | Within-region (stratified) | Default | - 0.283 | 0.114 | 0.030 |
|  |  |  | Within-region (stratified) | DorsAttn | - 0.013 | 0.113 | 0.912 |
|  |  |  | Within-region (stratified) | Limbic | - 0.404 | 0.220 | 0.094 |
|  |  |  | Within-region (stratified) | SalVentAttn | - 0.477 | 0.149 | 0.005 |
|  |  |  | Within-region (stratified) | SomMot | - 0.422 | 0.108 | <0.001 |
|  |  |  | Within-region (stratified) | Vis | - 0.299 | 0.127 | 0.032 |

#### 2.3.3. Relationship between synaptic terminal SV2A density and fractional anisotropy (FA)

Fractional anisotropy (FA) was used as a secondary marker of tissue microstructural organisation. Linear mixed models were fitted with synaptic density (BP_ND_ or V_T_) as the outcome and FA as predictor, using the same within-between model as for ODI (predictor split into a within-component and subject and region means, with random intercepts for subject and ROI). Results are reported in Supplementary Table 5.

**Supplementary Table 5.**
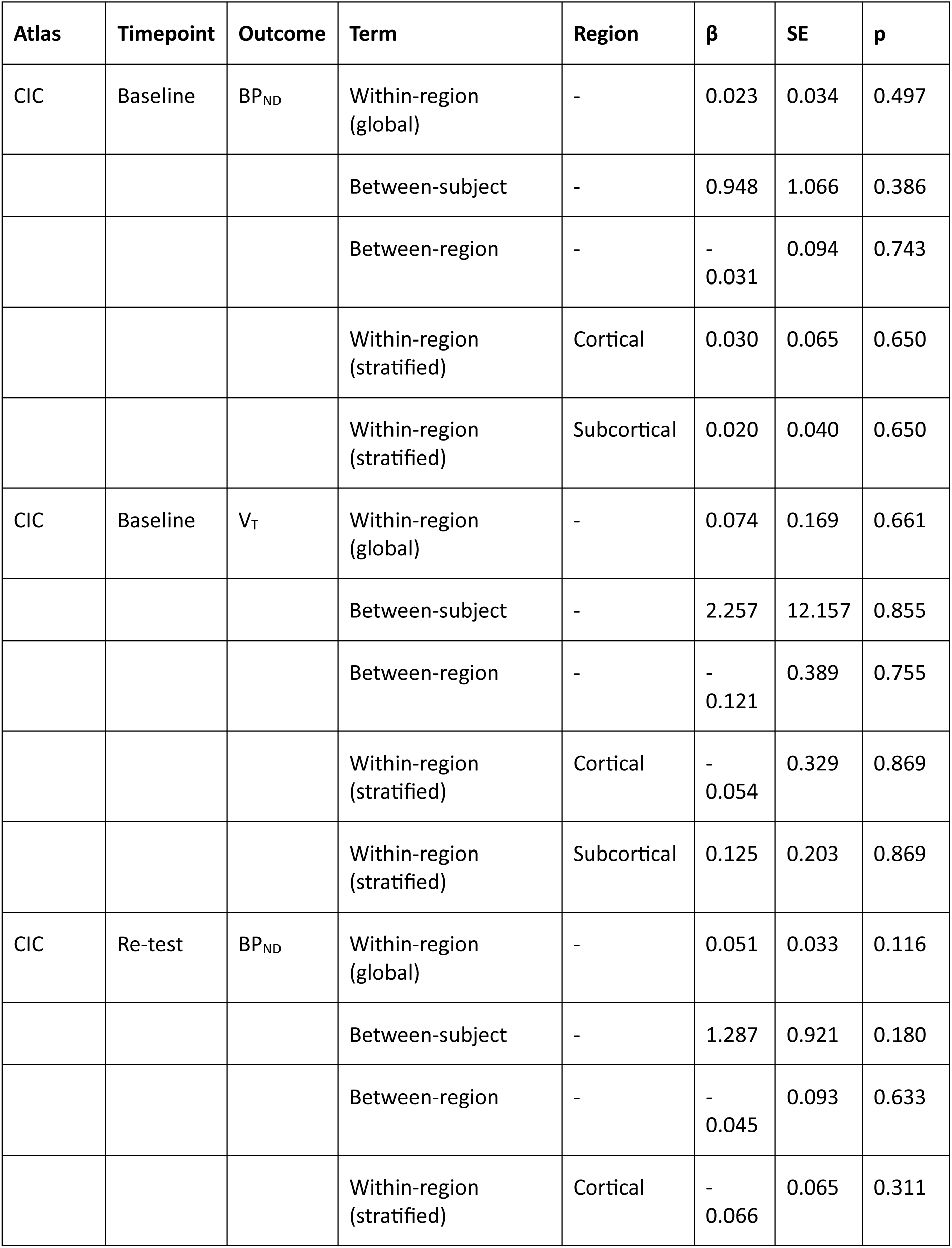
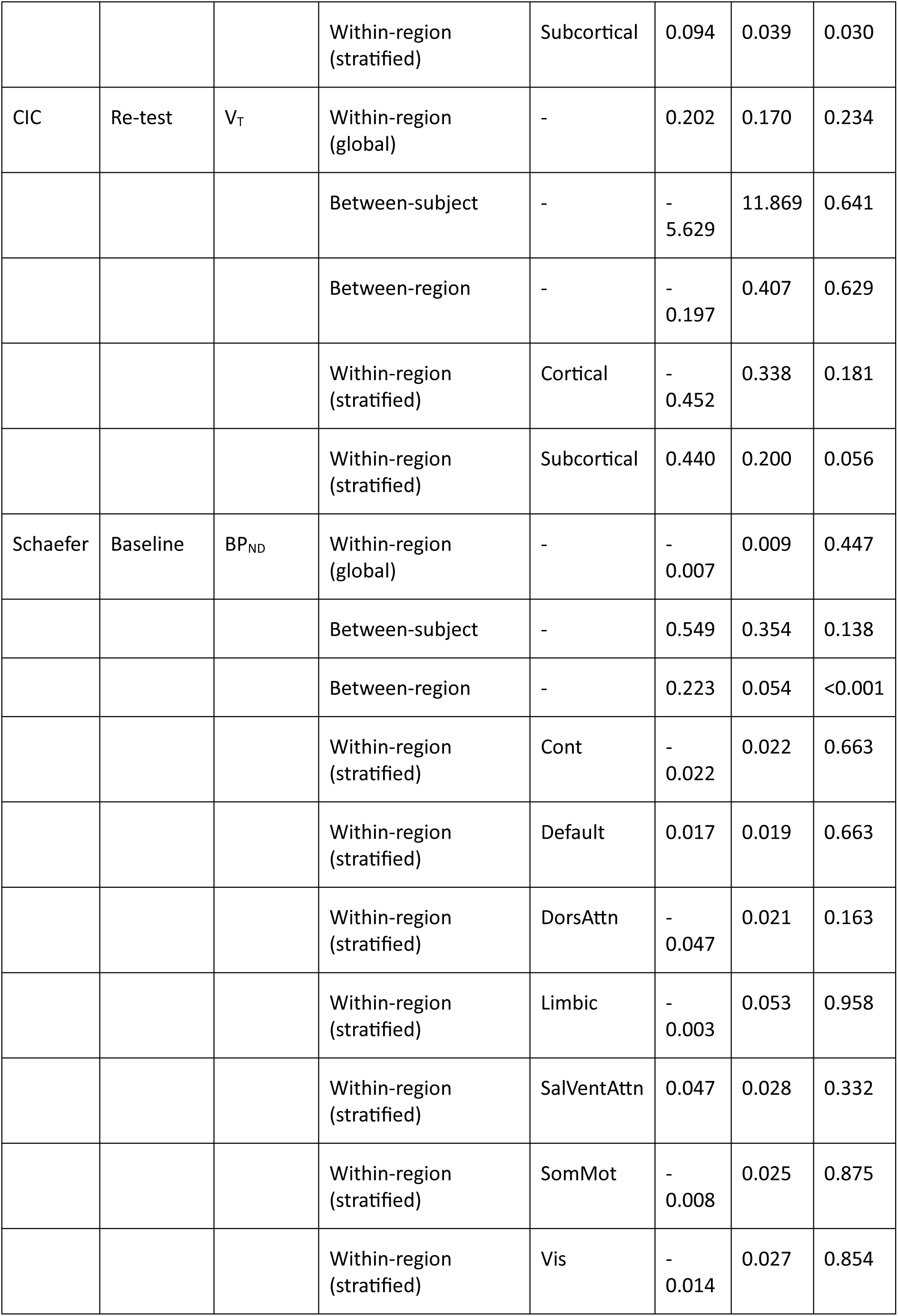
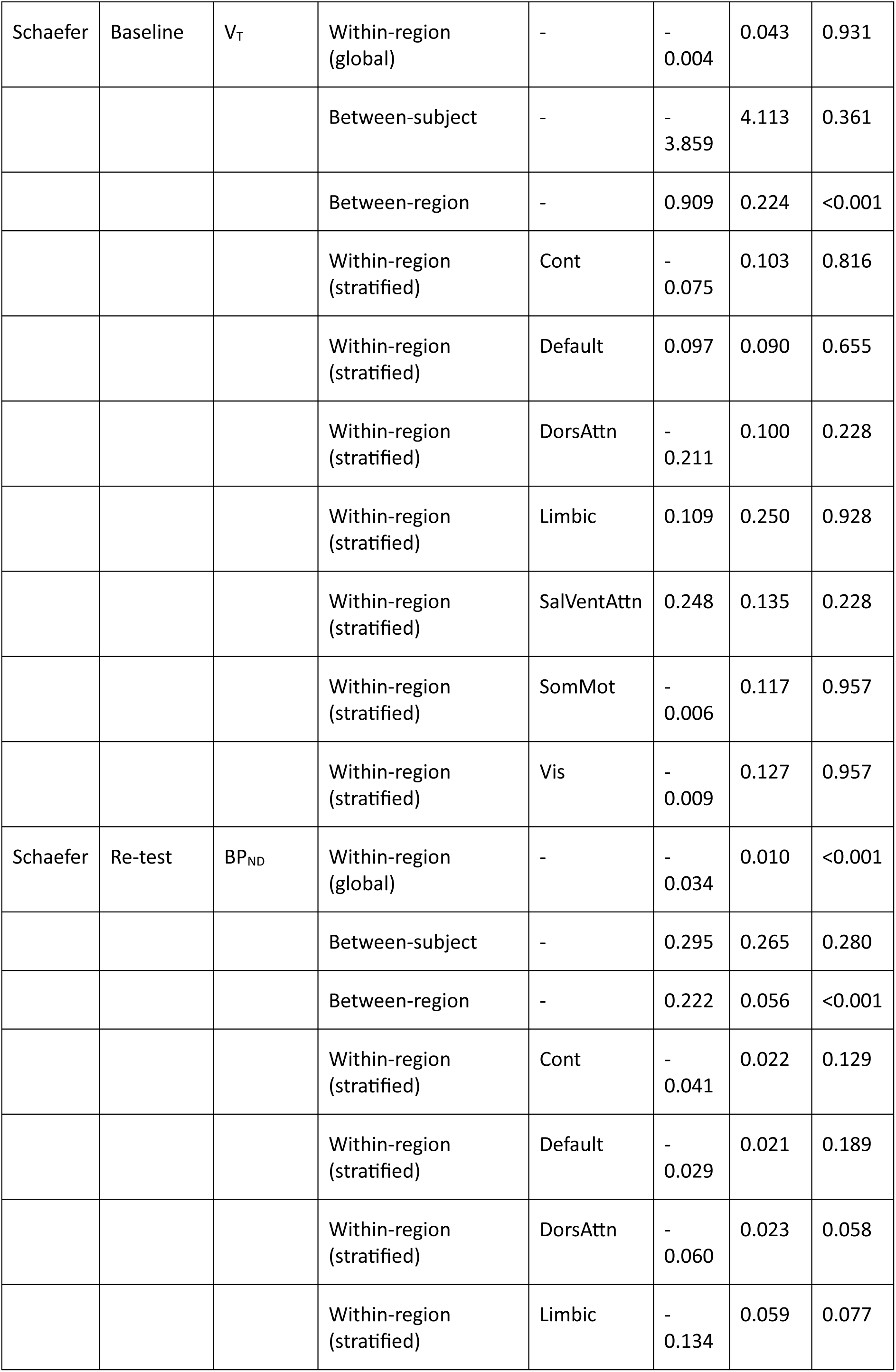
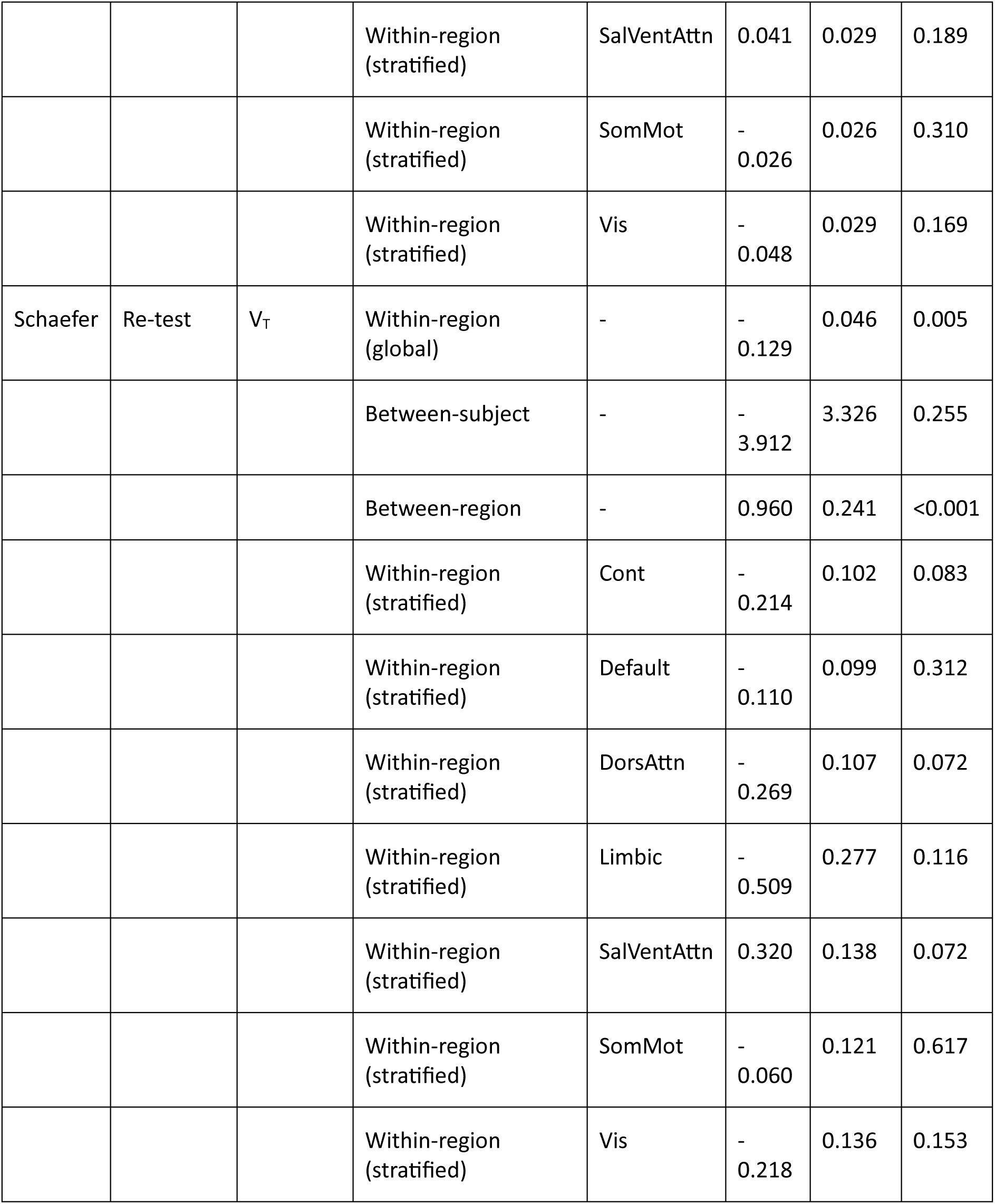
Associations between fractional anisotropy (FA) and synaptic terminal SV2A density (BP_ND_ and V_T_) in the CIC atlas (cortical and subcortical regions) and the Schaefer atlas (100 parcels, seven networks), at baseline and re-test. Within-between linear mixed models with random intercepts for subject and region. Within-region (global): within-region term of the global model (maximum likelihood); Between-subject and Between-region: subject-mean and region-mean terms of the same model; Within-region (stratified): cortical/subcortical or network-specific within-region slopes from the region-stratified model (restricted maximum likelihood; Kenward-Roger degrees of freedom), with p-values FDR-corrected across the subdivisions of each model. β is the change in BP_ND_ or V_T_ per pooled SD of FA; because subject means vary much less than individual values, between-subject β values are correspondingly large. Networks: Cont, control; Default, default mode; DorsAttn, dorsal attention; SalVentAttn, salience/ventral attention; SomMot, somatomotor; Vis, visual.

| Atlas | Timepoint | Outcome | Term | Region | $\beta$ | SE | p |
| --- | --- | --- | --- | --- | --- | --- | --- |
| CIC | Baseline | $BP_{ND}$ | Within-region (global) | - | 0.023 | 0.034 | 0.497 |
|  |  |  | Between-subject | - | 0.948 | 1.066 | 0.386 |
|  |  |  | Between-region | - | -0.031 | 0.094 | 0.743 |
|  |  |  | Within-region (stratified) | Cortical | 0.030 | 0.065 | 0.650 |
|  |  |  | Within-region (stratified) | Subcortical | 0.020 | 0.040 | 0.650 |
| CIC | Baseline | $V_T$ | Within-region (global) | - | 0.074 | 0.169 | 0.661 |
|  |  |  | Between-subject | - | 2.257 | 12.157 | 0.855 |
|  |  |  | Between-region | - | -0.121 | 0.389 | 0.755 |
|  |  |  | Within-region (stratified) | Cortical | -0.054 | 0.329 | 0.869 |
|  |  |  | Within-region (stratified) | Subcortical | 0.125 | 0.203 | 0.869 |
| CIC | Re-test | $BP_{ND}$ | Within-region (global) | - | 0.051 | 0.033 | 0.116 |
|  |  |  | Between-subject | - | 1.287 | 0.921 | 0.180 |
|  |  |  | Between-region | - | -0.045 | 0.093 | 0.633 |
|  |  |  | Within-region (stratified) | Cortical | -0.066 | 0.065 | 0.311 |
|  |  |  | Within-region (stratified) | Subcortical | 0.094 | 0.039 | 0.030 |
| CIC | Re-test | V<sub>T</sub> | Within-region (global) | - | 0.202 | 0.170 | 0.234 |
|  |  |  | Between-subject | - | - 5.629 | 11.869 | 0.641 |
|  |  |  | Between-region | - | - 0.197 | 0.407 | 0.629 |
|  |  |  | Within-region (stratified) | Cortical | - 0.452 | 0.338 | 0.181 |
|  |  |  | Within-region (stratified) | Subcortical | 0.440 | 0.200 | 0.056 |
| Schaefer | Baseline | BP<sub>ND</sub> | Within-region (global) | - | - 0.007 | 0.009 | 0.447 |
|  |  |  | Between-subject | - | 0.549 | 0.354 | 0.138 |
|  |  |  | Between-region | - | 0.223 | 0.054 | <0.001 |
|  |  |  | Within-region (stratified) | Cont | - 0.022 | 0.022 | 0.663 |
|  |  |  | Within-region (stratified) | Default | 0.017 | 0.019 | 0.663 |
|  |  |  | Within-region (stratified) | DorsAttn | - 0.047 | 0.021 | 0.163 |
|  |  |  | Within-region (stratified) | Limbic | - 0.003 | 0.053 | 0.958 |
|  |  |  | Within-region (stratified) | SalVentAttn | 0.047 | 0.028 | 0.332 |
|  |  |  | Within-region (stratified) | SomMot | - 0.008 | 0.025 | 0.875 |
|  |  |  | Within-region (stratified) | Vis | - 0.014 | 0.027 | 0.854 |
| Schaefer | Baseline | V<sub>T</sub> | Within-region (global) | - | - 0.004 | 0.043 | 0.931 |
|  |  |  | Between-subject | - | - 3.859 | 4.113 | 0.361 |
|  |  |  | Between-region | - | 0.909 | 0.224 | <0.001 |
|  |  |  | Within-region (stratified) | Cont | - 0.075 | 0.103 | 0.816 |
|  |  |  | Within-region (stratified) | Default | 0.097 | 0.090 | 0.655 |
|  |  |  | Within-region (stratified) | DorsAttn | - 0.211 | 0.100 | 0.228 |
|  |  |  | Within-region (stratified) | Limbic | 0.109 | 0.250 | 0.928 |
|  |  |  | Within-region (stratified) | SalVentAttn | 0.248 | 0.135 | 0.228 |
|  |  |  | Within-region (stratified) | SomMot | - 0.006 | 0.117 | 0.957 |
|  |  |  | Within-region (stratified) | Vis | - 0.009 | 0.127 | 0.957 |
| Schaefer | Re-test | BP<sub>ND</sub> | Within-region (global) | - | - 0.034 | 0.010 | <0.001 |
|  |  |  | Between-subject | - | 0.295 | 0.265 | 0.280 |
|  |  |  | Between-region | - | 0.222 | 0.056 | <0.001 |
|  |  |  | Within-region (stratified) | Cont | - 0.041 | 0.022 | 0.129 |
|  |  |  | Within-region (stratified) | Default | - 0.029 | 0.021 | 0.189 |
|  |  |  | Within-region (stratified) | DorsAttn | - 0.060 | 0.023 | 0.058 |
|  |  |  | Within-region (stratified) | Limbic | - 0.134 | 0.059 | 0.077 |
|  |  |  | Within-region (stratified) | SalVentAttn | 0.041 | 0.029 | 0.189 |
|  |  |  | Within-region (stratified) | SomMot | - 0.026 | 0.026 | 0.310 |
|  |  |  | Within-region (stratified) | Vis | - 0.048 | 0.029 | 0.169 |
| Schaefer | Re-test | $V_T$ | Within-region (global) | - | - 0.129 | 0.046 | 0.005 |
|  |  |  | Between-subject | - | - 3.912 | 3.326 | 0.255 |
|  |  |  | Between-region | - | 0.960 | 0.241 | <0.001 |
|  |  |  | Within-region (stratified) | Cont | - 0.214 | 0.102 | 0.083 |
|  |  |  | Within-region (stratified) | Default | - 0.110 | 0.099 | 0.312 |
|  |  |  | Within-region (stratified) | DorsAttn | - 0.269 | 0.107 | 0.072 |
|  |  |  | Within-region (stratified) | Limbic | - 0.509 | 0.277 | 0.116 |
|  |  |  | Within-region (stratified) | SalVentAttn | 0.320 | 0.138 | 0.072 |
|  |  |  | Within-region (stratified) | SomMot | - 0.060 | 0.121 | 0.617 |
|  |  |  | Within-region (stratified) | Vis | - 0.218 | 0.136 | 0.153 |

#### 2.3.4. Relationship between synaptic terminal SV2A density and structural connectivity

Structural connectivity weighted degree (SWD) was indexed by the weighted degree (node strength) of each ROI, derived from the DWI connectome tractography for CIC and Schaefer atlases (for CIC, restricted to connections with the 121 grey-matter regions). Linear mixed models were fitted with synaptic density PET markers (BP_ND_ or V_T_) as the outcome and SWD as the predictor, using the same within-between model as for ODI (predictor split into a within-component and subject and region means, with random intercepts for subject and ROI). Results are reported in Supplementary Table 6.

**Supplementary Table 6.** Associations between structural weighted degree (SWD) and synaptic terminal SV2A density (BP_ND_ and V_T_) in the CIC atlas (cortical and subcortical regions) and the Schaefer atlas (100 parcels, seven networks), at baseline and re-test. Within-between linear mixed models with random intercepts for subject and region. Within-region (global): within-region term of the global model (maximum likelihood); Between-subject and Between-region: subject-mean and region-mean terms of the same model; Within-region (stratified): cortical/subcortical or network-specific within-region slopes from the region-stratified model (restricted maximum likelihood; Kenward-Roger degrees of freedom), with p-values FDR-corrected across the subdivisions of each model. β is the change in BP_ND_ or V_T_ per pooled SD of SWD; because subject means vary much less than individual values, between-subject β values are correspondingly large. Networks: Cont, control; Default, default mode; DorsAttn, dorsal attention; SalVentAttn, salience/ventral attention; SomMot, somatomotor; Vis, visual.

| Atlas | Timepoint | Outcome | Term | Region | $\beta$ | SE | p |
| --- | --- | --- | --- | --- | --- | --- | --- |
| CIC | Baseline | $BP_{ND}$ | Within-region (global) | - | 0.102 | 0.039 | 0.009 |
|  |  |  | Between-subject | - | -3.410 | 1.969 | 0.101 |
|  |  |  | Between-region | - | 0.123 | 0.093 | 0.188 |
|  |  |  | Within-region (stratified) | Cortical | 0.076 | 0.046 | 0.097 |
|  |  |  | Within-region (stratified) | Subcortical | 0.179 | 0.078 | 0.046 |
| CIC | Baseline | $V_T$ | Within-region (global) | - | 0.338 | 0.197 | 0.087 |
|  |  |  | Between-subject | - | -37.666 | 22.066 | 0.105 |
|  |  |  | Between-region | - | 0.503 | 0.384 | 0.192 |
|  |  |  | Within-region (stratified) | Cortical | 0.241 | 0.230 | 0.296 |
|  |  |  | Within-region (stratified) | Subcortical | 0.619 | 0.396 | 0.236 |
| CIC | Re-test | $BP_{ND}$ | Within-region (global) | - | 0.069 | 0.042 | 0.101 |
|  |  |  | Between-subject | - | -1.928 | 1.563 | 0.234 |
|  |  |  | Between-region | - | 0.114 | 0.092 | 0.219 |
|  |  |  | Within-region (stratified) | Cortical | 0.064 | 0.049 | 0.328 |
|  |  |  | Within-region (stratified) | Subcortical | 0.086 | 0.088 | 0.328 |
| CIC | Re-test | V<sub>T</sub> | Within-region (global) | - | 0.206 | 0.219 | 0.346 |
|  |  |  | Between-subject | - | - 36.777 | 18.054 | 0.057 |
|  |  |  | Between-region | - | 0.504 | 0.401 | 0.212 |
|  |  |  | Within-region (stratified) | Cortical | 0.260 | 0.252 | 0.603 |
|  |  |  | Within-region (stratified) | Subcortical | 0.031 | 0.456 | 0.945 |
| Schaefer | Baseline | BP<sub>ND</sub> | Within-region (global) | - | 0.026 | 0.012 | 0.027 |
|  |  |  | Between-subject | - | -0.357 | 0.702 | 0.618 |
|  |  |  | Between-region | - | -0.120 | 0.052 | 0.025 |
|  |  |  | Within-region (stratified) | Cont | -0.020 | 0.038 | 0.769 |
|  |  |  | Within-region (stratified) | Default | 0.030 | 0.025 | 0.519 |
|  |  |  | Within-region (stratified) | DorsAttn | -0.015 | 0.035 | 0.769 |
|  |  |  | Within-region (stratified) | Limbic | 0.112 | 0.044 | 0.038 |
|  |  |  | Within-region (stratified) | SalVentAttn | -0.028 | 0.032 | 0.690 |
|  |  |  | Within-region (stratified) | SomMot | -0.006 | 0.034 | 0.861 |
|  |  |  | Within-region (stratified) | Vis | 0.079 | 0.024 | 0.007 |
| Schaefer | Baseline | V<sub>T</sub> | Within-region (global) | - | 0.117 | 0.056 | 0.036 |
|  |  |  | Between-subject | - | -16.286 | 6.898 | 0.030 |
|  |  |  | Between-region | - | -0.493 | 0.217 | 0.025 |
|  |  |  | Within-region (stratified) | Cont | -0.050 | 0.183 | 0.783 |
|  |  |  | Within-region (stratified) | Default | 0.227 | 0.119 | 0.131 |
|  |  |  | Within-region (stratified) | DorsAttn | -0.126 | 0.167 | 0.783 |
|  |  |  | Within-region (stratified) | Limbic | 0.597 | 0.210 | 0.031 |
|  |  |  | Within-region (stratified) | SalVentAttn | -0.052 | 0.154 | 0.783 |
|  |  |  | Within-region (stratified) | SomMot | -0.054 | 0.164 | 0.783 |
|  |  |  | Within-region (stratified) | Vis | 0.230 | 0.114 | 0.131 |
| Schaefer | Re-test | BP<sub>ND</sub> | Within-region (global) | - | 0.010 | 0.012 | 0.391 |
|  |  |  | Between-subject | - | -0.658 | 0.613 | 0.297 |
|  |  |  | Between-region | - | -0.125 | 0.053 | 0.021 |
|  |  |  | Within-region (stratified) | Cont | -0.059 | 0.036 | 0.368 |
|  |  |  | Within-region (stratified) | Default | 0.010 | 0.026 | 0.698 |
|  |  |  | Within-region (stratified) | DorsAttn | -0.033 | 0.032 | 0.446 |
|  |  |  | Within-region (stratified) | Limbic | 0.046 | 0.046 | 0.446 |
|  |  |  | Within-region (stratified) | SalVentAttn | -0.042 | 0.039 | 0.446 |
|  |  |  | Within-region (stratified) | SomMot | -0.025 | 0.039 | 0.607 |
|  |  |  | Within-region (stratified) | Vis | 0.095 | 0.025 | 0.001 |
| Schaefer | Re-test | $V_T$ | Within-region (global) | - | 0.090 | 0.057 | 0.117 |
|  |  |  | Between-subject | - | -16.497 | 6.950 | 0.029 |
|  |  |  | Between-region | - | -0.539 | 0.231 | 0.021 |
|  |  |  | Within-region (stratified) | Cont | -0.087 | 0.171 | 0.715 |
|  |  |  | Within-region (stratified) | Default | 0.230 | 0.121 | 0.203 |
|  |  |  | Within-region (stratified) | DorsAttn | -0.191 | 0.152 | 0.368 |
|  |  |  | Within-region (stratified) | Limbic | 0.296 | 0.219 | 0.368 |
|  |  |  | Within-region (stratified) | SalVentAttn | -0.047 | 0.183 | 0.795 |
|  |  |  | Within-region (stratified) | SomMot | -0.175 | 0.182 | 0.470 |
|  |  |  | Within-region (stratified) | Vis | 0.315 | 0.118 | 0.053 |

#### 2.3.5. Relationship between synaptic terminal SV2A density and functional connectivity

Functional weighted degree (FWD) was computed from resting-state fMRI as the row-wise mean of the Spearman rank correlation connectivity matrix, i.e., the average Spearman correlation between each region’s time series and those of all regions of the atlas, for both the CIC and Schaefer atlases. Linear mixed models were fitted with synaptic density (BP_ND_ or V_T_) as the outcome and FWD as the predictor, using the same within-between model as for ODI (predictor split into a within-component and subject and region means, with random intercepts for subject and ROI), and network-specific slopes were estimated as described above. Results are reported in Supplementary Table 7.

**Supplementary Table 7.** Associations between functional weighted degree (FWD) and synaptic terminal SV2A density (BP_ND_ and V_T_) in the CIC atlas (cortical and subcortical regions) and the Schaefer atlas (100 parcels, seven networks), at baseline and re-test. Within-between linear mixed models with random intercepts for subject and region. Within-region (global): within-region term of the global model (maximum likelihood); Between-subject and Between-region: subject-mean and region-mean terms of the same model; Within-region (stratified): cortical/subcortical or network-specific within-region slopes from the region-stratified model (restricted maximum likelihood; Kenward-Roger degrees of freedom), with p-values FDR-corrected across the subdivisions of each model. β is the change in BP_ND_ or V_T_ per pooled SD of FWD; because subject means vary much less than individual values, between-subject β values are correspondingly large. Networks: Cont, control; Default, default mode; DorsAttn, dorsal attention; SalVentAttn, salience/ventral attention; SomMot, somatomotor; Vis, visual.

| Atlas | Timepoint | Outcome | Term | Region | $\beta$ | SE | p |
| --- | --- | --- | --- | --- | --- | --- | --- |
| CIC | Baseline | $BP_{ND}$ | Within-region (global) | - | -0.012 | 0.010 | 0.240 |
|  |  |  | Between-subject | - | 0.187 | 0.120 | 0.136 |
|  |  |  | Between-region | - | 0.473 | 0.144 | 0.001 |
|  |  |  | Within-region (stratified) | Cortical | -0.041 | 0.014 | 0.009 |
|  |  |  | Within-region (stratified) | Subcortical | 0.023 | 0.016 | 0.143 |
| CIC | Baseline | $V_T$ | Within-region (global) | - | -0.086 | 0.052 | 0.103 |
|  |  |  | Between-subject | - | -1.765 | 1.372 | 0.215 |
|  |  |  | Between-region | - | 1.970 | 0.596 | 0.001 |
|  |  |  | Within-region (stratified) | Cortical | -0.200 | 0.072 | 0.011 |
|  |  |  | Within-region (stratified) | Subcortical | 0.058 | 0.081 | 0.475 |
| CIC | Re-test | $BP_{ND}$ | Within-region (global) | - | 0.022 | 0.013 | 0.087 |
|  |  |  | Between-subject | - | -0.054 | 0.093 | 0.567 |
|  |  |  | Between-region | - | 0.448 | 0.160 | 0.006 |
|  |  |  | Within-region (stratified) | Cortical | 0.030 | 0.018 | 0.207 |
|  |  |  | Within-region (stratified) | Subcortical | 0.014 | 0.019 | 0.461 |
| CIC | Re-test | V<sub>T</sub> | Within-region (global) | - | 0.085 | 0.066 | 0.201 |
|  |  |  | Between-subject | - | 0.436 | 1.153 | 0.710 |
|  |  |  | Between-region | - | 1.961 | 0.698 | 0.006 |
|  |  |  | Within-region (stratified) | Cortical | 0.096 | 0.094 | 0.452 |
|  |  |  | Within-region (stratified) | Subcortical | 0.073 | 0.097 | 0.452 |
| Schaefer | Baseline | BP<sub>ND</sub> | Within-region (global) | - | -<br>0.003 | 0.009 | 0.712 |
|  |  |  | Between-subject | - | 0.140 | 0.098 | 0.169 |
|  |  |  | Between-region | - | 0.154 | 0.131 | 0.241 |
|  |  |  | Within-region (stratified) | Cont | -<br>0.053 | 0.020 | 0.027 |
|  |  |  | Within-region (stratified) | Default | 0.038 | 0.017 | 0.063 |
|  |  |  | Within-region (stratified) | DorsAttn | -<br>0.042 | 0.027 | 0.209 |
|  |  |  | Within-region (stratified) | Limbic | -<br>0.028 | 0.031 | 0.430 |
|  |  |  | Within-region (stratified) | SalVentAttn | -<br>0.005 | 0.024 | 0.830 |
|  |  |  | Within-region (stratified) | SomMot | 0.081 | 0.027 | 0.022 |
|  |  |  | Within-region (stratified) | Vis | -<br>0.027 | 0.023 | 0.345 |
| Schaefer | Baseline | V<sub>T</sub> | Within-region (global) | - | 0.043 | 0.041 | 0.288 |
|  |  |  | Between-subject | - | -<br>0.977 | 1.133 | 0.400 |
|  |  |  | Between-region | - | 0.638 | 0.540 | 0.240 |
|  |  |  | Within-region<br>(stratified) | Cont | -<br>0.200 | 0.094 | 0.077 |
|  |  |  | Within-region<br>(stratified) | Default | 0.246 | 0.081 | 0.009 |
|  |  |  | Within-region<br>(stratified) | DorsAttn | -<br>0.178 | 0.128 | 0.230 |
|  |  |  | Within-region<br>(stratified) | Limbic | -<br>0.214 | 0.147 | 0.230 |
|  |  |  | Within-region<br>(stratified) | SalVentAttn | 0.059 | 0.114 | 0.603 |
|  |  |  | Within-region<br>(stratified) | SomMot | 0.589 | 0.129 | <0.001 |
|  |  |  | Within-region<br>(stratified) | Vis | -<br>0.101 | 0.111 | 0.421 |
| Schaefer | Re-test | BP<sub>ND</sub> | Within-region (global) | - | 0.005 | 0.011 | 0.612 |
|  |  |  | Between-subject | - | -<br>0.007 | 0.087 | 0.935 |
|  |  |  | Between-region | - | 0.109 | 0.139 | 0.437 |
|  |  |  | Within-region<br>(stratified) | Cont | -<br>0.040 | 0.023 | 0.188 |
|  |  |  | Within-region<br>(stratified) | Default | 0.015 | 0.021 | 0.544 |
|  |  |  | Within-region<br>(stratified) | DorsAttn | -<br>0.015 | 0.035 | 0.666 |
|  |  |  | Within-region<br>(stratified) | Limbic | 0.087 | 0.044 | 0.164 |
|  |  |  | Within-region<br>(stratified) | SalVentAttn | -<br>0.049 | 0.034 | 0.266 |
|  |  |  | Within-region (stratified) | SomMot | 0.065 | 0.032 | 0.164 |
|  |  |  | Within-region (stratified) | Vis | 0.028 | 0.030 | 0.486 |
| Schaefer | Re-test | V<sub>T</sub> | Within-region (global) | - | 0.058 | 0.051 | 0.253 |
|  |  |  | Between-subject | - | 0.476 | 1.091 | 0.668 |
|  |  |  | Between-region | - | 0.486 | 0.604 | 0.423 |
|  |  |  | Within-region (stratified) | Cont | -<br>0.124 | 0.109 | 0.449 |
|  |  |  | Within-region (stratified) | Default | 0.160 | 0.100 | 0.280 |
|  |  |  | Within-region (stratified) | DorsAttn | -<br>0.123 | 0.165 | 0.529 |
|  |  |  | Within-region (stratified) | Limbic | 0.322 | 0.207 | 0.280 |
|  |  |  | Within-region (stratified) | SalVentAttn | -<br>0.134 | 0.161 | 0.529 |
|  |  |  | Within-region (stratified) | SomMot | 0.372 | 0.149 | 0.090 |
|  |  |  | Within-region (stratified) | Vis | 0.041 | 0.140 | 0.769 |

#### 2.3.6. Relationship between synaptic terminal SV2A density and short-term plasticity

Short-term plasticity was indexed by the repetition suppression (RS) of the roving MMN, quantified for each participant as the slope of the MMN amplitude at Fz across tone positions S1-S6, computed from the mean amplitude (primary measure) and, as a sensitivity measure, from the peak amplitude. Pearson correlations (n = 18) were computed between the RS slope and the volume-weighted BP_ND_ or V_T_ of the frontotemporal ROI group, separately for the baseline and re-test scans and for the CIC and Schaefer atlases. As a control, the amplitude of the response to the deviant tone (S1) was correlated with frontotemporal BP_ND_ (Schaefer atlas, baseline). All results are summarised in Supplementary Table 8.

**Supplementary Table 8.**
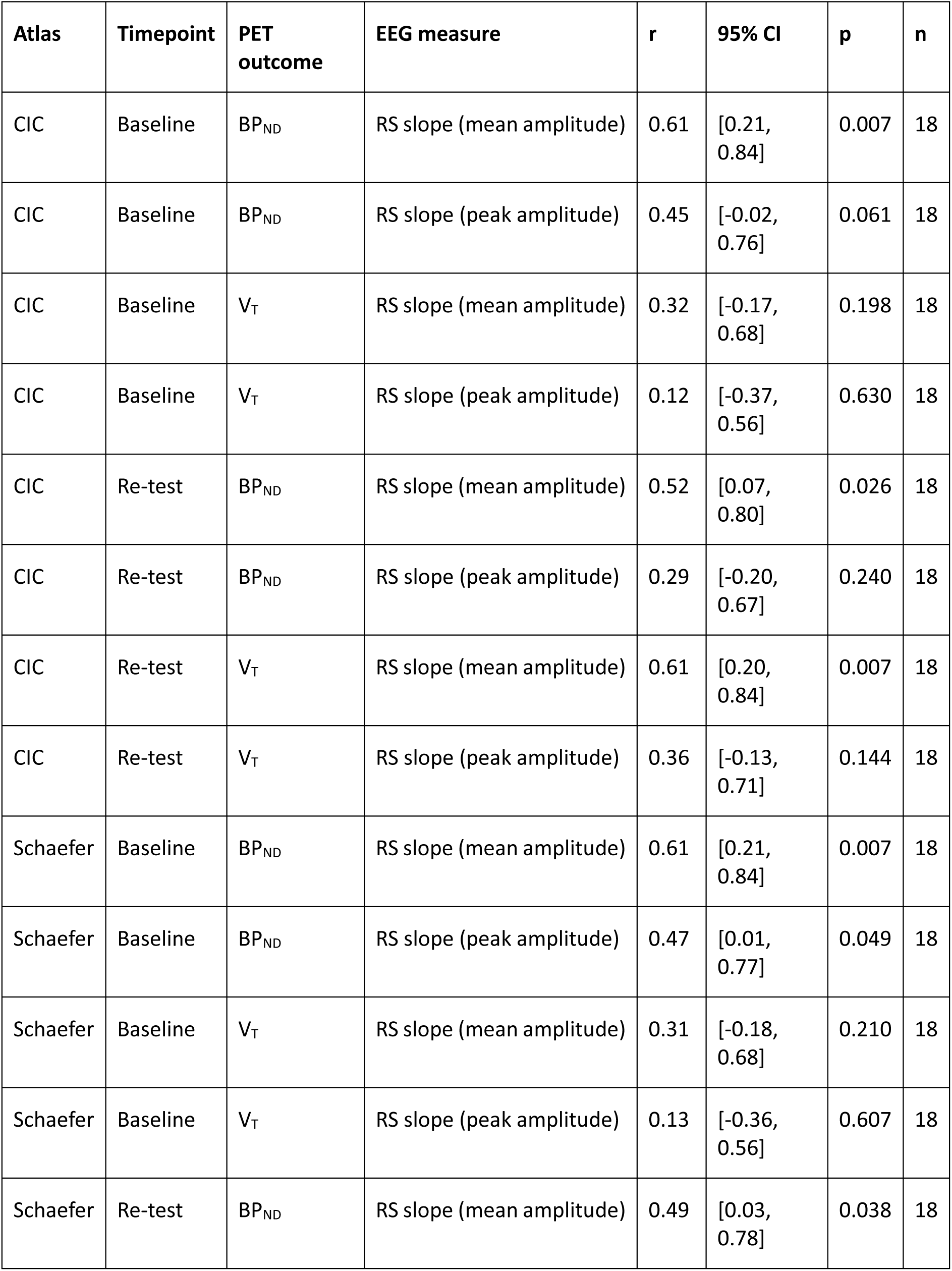
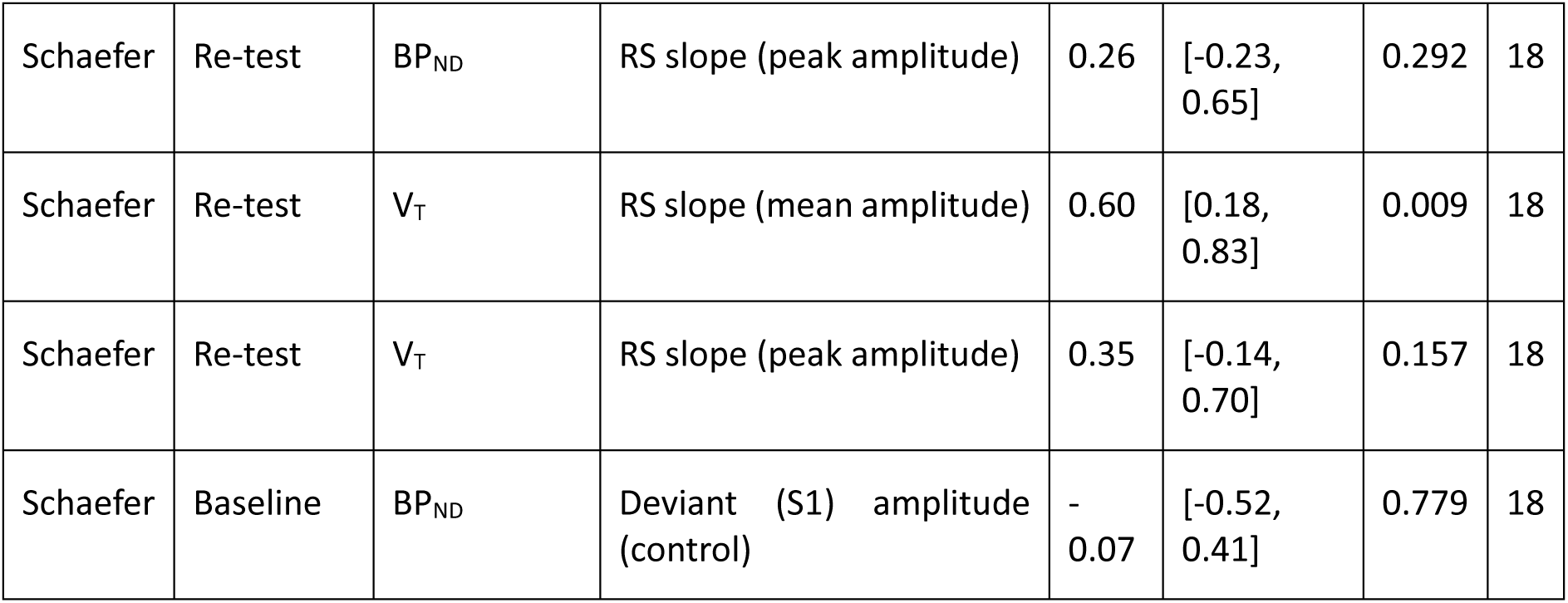
Pearson correlations between MMN repetition suppression (RS) and frontotemporal synaptic terminal SV2A density (volume-weighted BP_ND_ and V_T_ of the frontotemporal ROI group) in the CIC and Schaefer atlases, at baseline and re-test. RS slope: slope of the MMN amplitude at Fz across tone positions S1-S6, computed from the mean amplitude (primary measure) or the peak amplitude. Deviant (S1) amplitude: control analysis. CI, confidence interval.

#### 2.3.7. Relationship between synaptic terminal SV2A density and Hebbian-like long-term plasticity

Hebbian-like long-term plasticity was indexed by late-phase P2 potentiation (ΔP2-Late), the difference in P2 peak amplitude at POz between the late post-tetanic and the pre-tetanus blocks of the vLTP task. Pearson correlations (n = 18) were computed between ΔP2-Late and the volume-weighted BP_ND_ or V_T_ of the occipital ROI group, for the CIC and Schaefer atlases. Correlations were tested at baseline only, because P2 potentiation was not significant at re-test (late vs pre-tetanus P2 peak amplitude: β = 0.580, p = 0.326; main manuscript); early-phase N1b potentiation was not tested because the N1b was not potentiated at baseline (β = -0.248, p = 0.154). As a control, the pre-tetanus P2 peak amplitude was correlated with occipital BP_ND_ (Schaefer atlas, baseline). All results are summarised in Supplementary Table 9.

**Supplementary Table 9.** Pearson correlations between late-phase P2 potentiation (ΔP2-Late: late post-tetanic minus pre-tetanus P2 peak amplitude at POz) and occipital synaptic terminal SV2A density (volume-weighted BP_ND_ and V_T_ of the occipital ROI group) in the CIC and Schaefer atlases at baseline. Pre-tetanus P2 peak amplitude: control analysis. CI, confidence interval.

| Atlas | Timepoint | PET outcome | EEG measure | r | 95% CI | p | n |
| --- | --- | --- | --- | --- | --- | --- | --- |
| CIC | Baseline | BP<sub>ND</sub> | ΔP2-Late | 0.53 | [0.09, 0.80] | 0.023 | 18 |
| CIC | Baseline | V<sub>T</sub> | ΔP2-Late | 0.68 | [0.31, 0.87] | 0.002 | 18 |
| Schaefer | Baseline | BP<sub>ND</sub> | ΔP2-Late | 0.54 | [0.09, 0.80] | 0.022 | 18 |
| Schaefer | Baseline | V<sub>T</sub> | ΔP2-Late | 0.67 | [0.29, 0.87] | 0.002 | 18 |
| Schaefer | Baseline | BP<sub>ND</sub> | Pre-tetanus P2 peak (control) | 0.11 | [-0.38, 0.55] | 0.676 | 18 |

#### 2.3.8. Synaptic terminal SV2A density, EEG plasticity markers and neurite complexity

To test whether the associations between the EEG plasticity markers and synaptic terminal SV2A density were independent of neurite complexity, each EEG marker was added as a covariate to the within-between ODI-BP_ND_ model, fitted at baseline to the regions of the ROI group used for the corresponding EEG-PET correlation (frontotemporal for MMN repetition suppression; occipital for late-phase P2 potentiation): BP_ND_ ∼ ODI_within_ + ODI_subject_ + ODI_region_ + EEG + (1 | Subject_ID) + (1 | ROI_Label), fitted by maximum likelihood. ODI was decomposed within the ROI group, and the EEG marker (one value per subject) was z-scored, so that its coefficient is the change in BP_ND_ per standard deviation of the EEG marker. MMN repetition suppression remained associated with frontotemporal BP_ND_ after adjustment for ODI in both the Schaefer (β = 0.210, p = 0.003) and CIC atlases (β = 0.186, p < 0.001), as did late-phase P2 potentiation with occipital BP_ND_ (Schaefer: β = 0.199, p = 0.023; CIC: β = 0.238, p = 0.010). Results are reported in Supplementary Table 10.

**Supplementary Table 10.** Associations between the EEG plasticity markers and synaptic terminal SV2A density (BP_ND_) after adjustment for neurite complexity (ODI), at baseline. Linear mixed models fitted by maximum likelihood to the regions of the ROI group used for each EEG-PET correlation (frontotemporal: 43 Schaefer parcels, 22 CIC regions; occipital: 17 Schaefer parcels, 12 CIC regions): BP_ND_ ∼ ODI_within_ + ODI_subject_ + ODI_region_ + EEG + (1 | Subject_ID) + (1 | ROI_Label). ODI terms: change in BP_ND_ per pooled SD of ODI (decomposed within the ROI group); EEG marker: change in BP_ND_ per SD of the EEG marker (z-scored). RS, repetition suppression; ΔP2-Late, late-phase P2 potentiation.

| Atlas | EEG marker (ROI group) | Term | $\beta$ | SE | p |
| --- | --- | --- | --- | --- | --- |
| Schaefer | MMN RS slope (frontotemporal) | ODI within-region | 0.021 | 0.012 | 0.073 |
| Schaefer | MMN RS slope (frontotemporal) | ODI between-subject | 0.271 | 0.250 | 0.293 |
| Schaefer | MMN RS slope (frontotemporal) | ODI between-region | -0.028 | 0.092 | 0.761 |
| Schaefer | MMN RS slope (frontotemporal) | EEG marker | 0.210 | 0.061 | 0.003 |
| Schaefer | $\Delta P2$-Late (occipital) | ODI within-region | 0.029 | 0.017 | 0.087 |
| Schaefer | $\Delta P2$-Late (occipital) | ODI between-subject | 0.133 | 0.225 | 0.561 |
| Schaefer | $\Delta P2$-Late (occipital) | ODI between-region | 0.271 | 0.132 | 0.056 |
| Schaefer | $\Delta P2$-Late (occipital) | EEG marker | 0.199 | 0.080 | 0.023 |
| CIC | MMN RS slope (frontotemporal) | ODI within-region | -0.079 | 0.045 | 0.079 |
| CIC | MMN RS slope (frontotemporal) | ODI between-subject | 1.404 | 0.413 | 0.003 |
| CIC | MMN RS slope (frontotemporal) | ODI between-region | 0.965 | 0.107 | <0.001 |
| CIC | MMN RS slope (frontotemporal) | EEG marker | 0.186 | 0.041 | <0.001 |
| CIC | $\Delta P2$-Late (occipital) | ODI within-region | 0.000 | 0.031 | 0.987 |
| CIC | $\Delta P2$-Late (occipital) | ODI between-subject | 0.501 | 0.414 | 0.242 |
| CIC | $\Delta P2$-Late (occipital) | ODI between-region | 0.275 | 0.147 | 0.085 |
| CIC | $\Delta P2$-Late (occipital) | EEG marker | 0.238 | 0.082 | 0.010 |

### 2.4. Test-retest reliability

Test-retest reliability between the baseline and the post-drug re-test scans was assessed in the Schaefer atlas. ICC(A,1) (two-way model, absolute agreement, single measure) was computed on one value per subject and scan, with region-level measures averaged across the 100 parcels or across the ROI group (Supplementary Table 11). Reliability was excellent for SWD (ICC = 0.91), good for ODI (ICC = 0.86), moderate for BP_ND_ (100 parcels: ICC = 0.70; frontotemporal: ICC = 0.71; occipital: ICC = 0.74) and the MMN repetition suppression slope (ICC = 0.56), and poor for ΔP2-Late (ICC = 0.02). The stability of the region-level associations was tested by fitting both scans together, with the predictor decomposed within each scan, and testing the interaction between the within-region term and scan: Outcome ∼ (X_within_ + X_subject_ + X_region_) × Scan + (1 | Subject_ID/Scan) + (1 | ROI_Label). The within-region ODI-BP_ND_ (change in β = 0.008, p = 0.459) and SWD-BP_ND_ (change in β = -0.016, p = 0.354) slopes did not differ between scans. The stability of the subject-level EEG-PET correlations was assessed with Zou’s 95% confidence interval for the difference between two dependent, non-overlapping correlations: neither the MMN repetition suppression-frontotemporal BP_ND_ correlation (Δr = -0.12, 95% CI -0.57 to 0.30) nor the ΔP2-Late-occipital BP_ND_ correlation (Δr = -0.37, 95% CI -0.93 to 0.24) changed significantly between scans. Results are reported in Supplementary Table 12.

**Supplementary Table 11.**
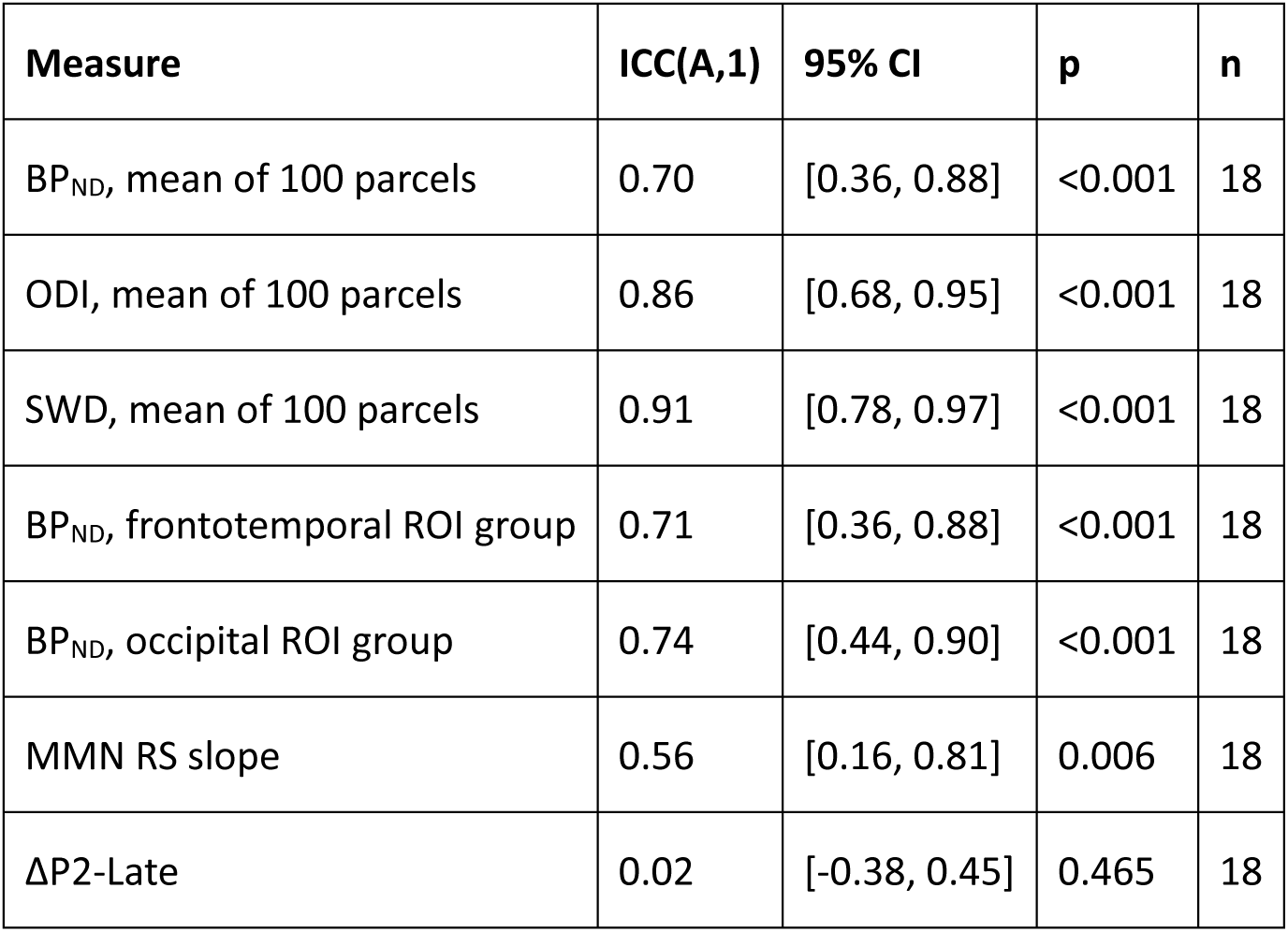
Test-retest reliability of the PET, MRI and EEG measures between the baseline and post-drug re-test scans (Schaefer atlas; n = 18). ICC(A,1): intraclass correlation coefficient, two-way model, absolute agreement, single measure. RS, repetition suppression; ΔP2-Late, late-phase P2 potentiation; CI, confidence interval.

**Supplementary Table 12.** Stability of the PET-MRI and EEG-PET associations between the baseline and post-drug re-test scans (Schaefer atlas; n = 18). Region-level slopes: within-region coefficient at baseline and its change at re-test (X_within_ × Scan interaction) from Outcome ∼ (X_within_ + X_subject_ + X_region_) × Scan + (1 | Subject_ID/Scan) + (1 | ROI_Label), with the predictor decomposed within each scan; β per pooled SD of the predictor. Subject-level correlations: Pearson’s r at baseline and the re-test minus baseline difference (Δr), with Zou’s 95% confidence interval for the difference between two dependent, non-overlapping correlations (an interval including zero indicates no significant change). RS, repetition suppression; ΔP2-Late, late-phase P2 potentiation; CI, confidence interval.

| Association | Baseline | Change at re-test | SE / 95% CI | p |
| --- | --- | --- | --- | --- |
| ODI-BP<sub>ND</sub> within-region slope ($\beta$) | 0.036 | 0.008 | SE = 0.011 | 0.459 |
| SWD-BP<sub>ND</sub> within-region slope ($\beta$) | 0.026 | -0.016 | SE = 0.017 | 0.354 |
| MMN RS slope - frontotemporal BP<sub>ND</sub> (r) | 0.61 | -0.12 | [-0.57, 0.30] | - |
| ΔP2-Late - occipital BP<sub>ND</sub> (r) | 0.54 | -0.37 | [-0.93, 0.24] | - |

